# Rye stem rust is a reservoir of diversity and virulence for wheat stem rust

**DOI:** 10.64898/2026.10.12.758526

**Authors:** Thomas W. Ingram, Molly L. Veregge, Pablo D. Olivera Firpo, Yue Jin, Alina Akhunova, Eduard Akhunov, Matthew N. Rouse, Matthew J. Moscou

**Affiliations:** Department of Plant Pathology, University of Minnesota, St. Paul, MN 55108, USA; USDA-ARS Cereal Disease Laboratory, University of Minnesota, St. Paul, MN 55108, USA; Integrated Genomics Facility, Kansas State University, Manhattan, KS 66506, USA; Department of Plant Pathology, Kansas State University, Manhattan, KS 66506, USA; USDA-ARS Sugarcane Field Station, Canal Point, FL 33438, USA

**Keywords:** *Puccinia graminis*, wheat, rye, stem rust, mating genes, admixture

## Abstract

Sexual populations of the stem rust pathogen (*Puccinia graminis* or *Pg*) are producing new race types globally. An analysis of 385 *Pg* genomes (204 from this study) illuminated international migration and admixture events.

Within Kazakhstan, a novel and diverse sexual clade of *P. graminis* is described. An analysis of the highly divergent mating type homeodomain (HD) locus found that the 75 Kazakhstan isolates had 41 different HD haplotypes, out of a total of 99 *P. graminis* HD haplotypes found globally. Kazakh isolates share HD haplotypes with isolates in clade 1 (Ug99), 3-B (TTRTF), and 4 (TKTTF).

There are 148 unique combinations of HD locus haplotypes within the 385 isolates. Rye (*Pgs*) and wheat (*Pgt*) stem rust share multiple HD locus haplotypes and have conserved pheromone receptors and precursors, while oat stem rust (*Pga*) does not.

Virulent alleles for *AvrSr13*, *AvrSr22*, and *AvrSr50* in *Pgt* are frequently derived from admixture with *Pgs*. *Pgs* also contains virulent effectors for *AvrSr33*, *AvrSrC2*, and *AvrSr27*. *Pgt* isolates virulent on *Sr31*, such as Ug99 and isolates from Spain, have genetic links to North American *Pgs* isolates which are present in regions of the United States where barberry is common.

## Introduction

*Puccinia graminis* (*Pg*) is a wind-borne basidiomycete fungal species that causes stem rust on cereal crops such as wheat, barley, rye, oats, and some wild grasses (Fetch et al., 2021). Specialized forms (*formae speciales* or ff. spp.) principally infect their corresponding host such as *P. graminis* f. sp. *tritici* (*Pgt*) on wheat (*Triticum aestivum*; *Triticum turgidum*) and barley (*Hordeum vulgare*), *P. graminis* f. sp. *secalis* (*Pgs*) on rye (*Secale cereale*) and barley, and *P. graminis* f. sp. *avenae* (*Pga*) on oats (*Avena sativa*) (Stakman et al., 1962; Sun & Steffenson, 2005). Stem rust can cause yield losses up to 100% in heavily infested fields and remains a global threat to grain production (Olivera et al., 2015). Due to the low profit margins of cereal production, and limited access to chemical protection in some regions, host resistance remains one of the most economically viable methods of disease management. Host resistance in cereals against biotrophs is typically conferred by single dominant major-effect resistance genes, often encoding nucleotide-binding site–leucine-rich repeat (NLR) proteins that recognize secreted pathogen effectors (avirulence proteins or Avr) (Dodds & Rathjen, 2010). Deployment of host resistance as single major-effect genes is often overcome by new strains of the pathogens soon after their release (Ngou et al., 2022). Despite significant breeding efforts, there is an ongoing emergence of highly virulent *Pgt* races such as TTTTF (isolated as 01MN84-A-1-2), TRTTF (06YEM34-1), TTRTF (14GEO189-1), TKTTF (13ETH18-1), and TTKTT (14KEN58-1) (Jin, 2005; Olivera et al., 2012; Olivera et al., 2015; Olivera et al., 2019; Patpour et al., 2016; Roelfs & Martens, 1988). *Pgt* race TTKTT, a highly virulent variant of the Ug99 (TTKSK) race group, was detected in Kenya in 2014 (Newcomb et al. 2016), appeared in Iraq in 2021 (Nazari et al., 2021), and reached Nepal in 2023 (Patpour et al., 2024). Previously undefeated resistance genes such as *Sr47*, *Sr32*, and *Sr40* have been rendered ineffective by sexually derived isolates in Kazakhstan (Olivera et al., 2022). Another population of concern identified in Spain contains isolates virulent on the *Sr31* gene and represent the first case of a *Pgt* isolate detected outside of the Ug99 lineage to have virulence on *Sr31* (Olivera et al., 2022; Veregge et al. 2024).

*Pg* has five morphological spore classes, three on grasses and two on alternate host plant species, such as common barberry (*Berberis vulgaris*) (Leonard and Szabo, 2005). In the spring, overwintering telia on decaying grasses undergo meiosis to produce haploid basidiospores which infect the alternate host, forming pycnia and haploid pycniospores. The fusion of haploid pycniospores with receptive pycnia hyphae forms dikaryotic aeciospores on the underside of the leaf, and these sexually derived aeciospores then infect cereal crops. Within cereal fields, *Pg* reproduces clonally via asexual dikaryotic urediniospores to create secondary infections (Roelfs, 1985). In the United States, an annual enphytotic *Pg* initiates from asexual populations from the southern United States and Mexico where *Pg* is able to survive the winter months. In 2024, and the past decade, the predominant *Pgt* race in central USA has been QFCSC (Kolmer et al. 2007; Zegeye et al., 2023). However, sexual populations from wild plants in the Berberidaceae family and in the Pacific North-West (PNW) are creating novel physiological races, threatening a relatively stable wheat industry in the USA (Jin et al., 2014; Upadhaya, GC Upadhaya, & Brueggeman, 2024). Sexual isolates and populations have also been reported in the United Kingdom, Georgia, Spain, Sweden, and Kazakhstan within the past decade (Lewis et al., 2018; Olivera et al., 2019; Olivera et al., 2022; Rodriguez-Algaba et al., 2024; Tsushima et al., 2022). Although somatic hybridization appeared to be involved in the creation of some important new strains such as Ug99 (Guo et al., 2022), admixture events remain the most common source of novel race types of *Pg* (Lewis et al., 2018; Olivera et al., 2019; Olivera et al., 2022; Rodriguez-Algaba et al., 2024; Tsushima et al., 2022).

Between 2015 and 2019, severe wheat stem rust outbreaks were observed in western Siberia and north Kazakhstan (Olivera et al., 2022; Shamanin et al., 2020). Systematic phenotyping of isolates from this epidemic identified 112 races that were genetically distinct from previously studied *Pgt* isolates (Olivera et al., 2022). Three previously universally resistant *R* genes, *Sr32* + *Sr40* and *Sr47*, became ineffective to emerged *Pgt* races RHHTM and RCMTM, respectively. The proposed hypothesis for the sudden breakdown of dozens of R genes is the presence of barberry in this region, which allowed *Pgt* isolates to recombine and create novel race types (Olivera et al., 2022). Barberry is native to both Siberia and central Asia, as well as the Tianshan mountain range which spans the Himalayas, northern Pakistan, and Southeast Kazakhstan (Tudi et al., 2022).

Sexuality in basidiomycetes is well understood, since many mating type genes have been cloned and experimentally validated (James, 2023). Rusts have a tetrapolar heterothallic mating system which encourages sexual recombination between genetically dissimilar individuals. Recent progress has been made in the *Puccinia* genus, with the pre-gamete fusion pheromone, pheromone receptor (PR), and post-gamete fusion homeodomain (HD) loci being described in *P. graminis* f. sp. *tritici*, *P. coronata* f. sp. *avenae, P. triticina*, and *P. striiformis* f. sp. *tritici* (Cuomo et al., 2017; Holden et al., 2023; Li et al., 2019; Luo et al., 2024). For post-gamete fusion mating to be successful, haploid pycniospores produced on the alternate host must fuse with receptive hyphae containing a different homeodomain (Holden et al., 2023; Roelfs, 1985). HD1 in *Puccinia* spp. is hypothesized to contain a DNA binding domain that activates a host of genes necessary for hyphal cell division and clamp connections formation. HD2 proteins from different HD haplotypes bind to HD1 proteins and shuttle them into one of the nuclei in the dikaryotic cell. Thus, the homeodomain possesses higher than normal levels of non-synonymous changes (Kiran et al., 2021). Conversely, the pheromone precursor (*Mfa*) and pheromone receptor (*Pra* alleles *STE3.2–2* and *STE3.2–3*) loci are under purifying selection pressure to ensure gametic fusion occurs in all HD haplotype combinations (Li et al., 2019; Luo et al., 2024). Given that mating type genes are expressed in wheat tissue during asexual growth, it is possible there is some level of genetic exchange happening during the clonal urediniospore stage, such as somatic hybridization which is known to happen in *Puccinia* spp. (Li et al., 2019; Luo et al., 2024).

Recombination plays a major role in the formation of novel races in *P. graminis*. This study aims to understand how cryptic genetic variation is contributing to new races and epidemics. Selection pressure will inevitably drive populations to accumulate virulence phenotypes, and admixture creates diversity of allelic variation in *P. graminis* which allows for the accumulation of virulence combinations that are capable of overcoming most of the resistance genes breeders can incorporate in a timely manner into high yielding cereal cultivars. Using comparative genomics on a global *Pg* population, we show that past admixture events between wheat and rye stem rust pathogens, and cryptic populations of *P. graminis* are contributing to stem rust pandemics.

## Materials and Methods

### Isolate increase and race confirmation

A total of 204 *Pg* isolates were increased in Biosafety Level 2 and Biosafety Level 3 facilities at the USDA-ARS Cereal Disease Laboratory and the University of Minnesota, respectively. Each isolate was increased on the susceptible wheat cultivar McNair 701, which was planted at a density of 30-35 plants per pot. Following germination, seedlings were treated with maleic hydrazide to suppress growth and enhance susceptibility to infection. Seven days after planting, individual pots were sprayed with urediniospores from single-pustule isolate suspended in Soltrol 170 oil (Chevron Phillips Chemical Company, The Woodlands, TX). Inoculated plants were incubated overnight in a humidity chamber in dark conditions and then transferred to a greenhouse maintained at 22-24°C. To prevent contamination, seedlings were covered by cellophane bags five days after inoculation. Urediniospores were collected two weeks post inoculation in a biosafety cabinet, desiccated for 7 days, and then stored in a −80°C freezer until DNA extraction. In addition, isolate purity and race identity were confirmed by inoculating each isolate onto the standard set of 20 differential lines used for *Pgt* race identification (Jin et al., 2008).

### DNA extraction for Illumina short read sequencing

All 204 *Pg* isolates had DNA extracted from desiccated urediniospores using the OmniPrep™ (G-Biosciences, St. Louis, MO, USA) protocol. These isolates originated from multiple locations; 57 North American isolates (all from the USA), 103 Asian isolates, 24 European isolates, and 20 African isolates. The remaining isolates were sequenced similarly in separate studies under project accession numbers PRJEB47693 (Tsushima et al. 2022), PRJNA803546 (Guo et al. 2022), and PRJNA952228 (Upadhaya et al. 2024). Combining isolates sequenced in this study and previously sequenced isolates there are a total of 111 Asian, 43 African, 203 North American, 1 South American, and 27 European isolates. For DNA extraction, 25 mg of diatomaceous earth and 25 mg of frozen desiccated spores were added to a 2 mL tube. The tube was shaken at speed 4 for 20 s using a MP FastPrep-24 (MP Biomedicals, Irvine, CA, USA). 600 µl of OmniPrep™ Genomic Lysis Buffer Mix was added to the tube and mixed by inversion. 6 µl of Longlife™ Proteinase K (G-Biosciences, St. Louis, MO, USA) was added to the tube, incubated at 60°C for 1 hour, mixed by inversion every 15 minutes. Tubes were cooled to room temperature (20-22°C), and 200 µl of chloroform was added, mixed by inversion, and spun in a centrifuge for 10 min at 14,000 rpm. The supernatant was transferred to a new tube, and 50 µl DNA stripping solution was added. The solution was mixed by inversion and incubated at 60°C for 10 min, then placed on ice for 5 min. 100 µl of precipitation solution was added and mixed by inversion, spun for 5 min at 14,000 rpm. The supernatant was added to a clean tube, and 500 µl of isopropanol was used for DNA precipitation. The tube was mixed by inversion and spun for 10 min at 14,000 rpm. The supernatant was discarded and washed with 700 µl of 70% ethanol, spun for 5 min at 14,000 rpm, this was repeated once before air drying the pellet. The pellet was resuspended in 100 µl of TE buffer. DNA quantity and quality was checked using a NanoDrop 1000 Spectrophotometer (Thermo Fisher Scientific, Waltham, MA, USA). High quality DNA was sent to the Integrated Genomics Facility, Kansas State University, Manhattan, KS, USA for 150 bp PE Illumina sequencing. An initial set of 80 isolates was sequenced using this approach. A total of 124 isolates were sequenced in a similar manner but with different Illumina (San Diego, CA, USA) short read sequencing technologies (Dataset S1).

### Phylogenomic analysis

Sequences from Olivera et al. (2019), Upadhaya et al. (2024), and Guo et al. (2022) along with 204 whole genome sequencing *Pg* data in this study were mapped to the 99KS76A reference genome (GCA_002762355.2 Guo et al. 2022) using bwa-mem (bwa/0.7.17). Bcftools (v1.2) mpileup was used to call variants from the resulting bam file with a setting of –variants only. The variant call file (VCF) was sorted using bcftools and filtered again to exclude reads with the following command -e ‘QUAL < 30 || DP < 3 || AF < 0.05’. The VCFs were zipped, indexed, and merged with bcftools merge −0 to impute reads. After vcf files were merged, they were filtered a second time excluding ‘QUAL < 36 || DP < 4 || AF < 0.10’, sorted, zipped, and indexed. The merged VCF was loaded into Tassel (v5.2.93), a neighbor-joining tree and a principal component analysis was performed (PC=3). Plotly and pandas were used to visualize the PCA (Chen et al., 2022; Wickham, 2011). T-BAS v2.3 (Carbone et al. 2019) was used to visualize the mitochondrial tree was developed similarly to the whole genome tree, using bwa-mem default parameters, and the reference genome used with the Ug99 reference genome contig VDEP01000353.1 that represents the mitochondrial genome for that isolate. Trees were displayed with iTOL v6 (Letunic & Bork, 2024). PLINK2 (v2.0.0) was used to linkage disequilibrium prune the whole genome sequencing derived SNP data set using parameters “--indep-pairwise 50 10 0.1”. For the complete data set (348 isolates), the total number of SNPs were reduced to 439,775, whereas for the reduced data set (62 isolates), the total number of SNPs were reduced to 633,816. Maximum phylogenetic analysis was performed using RAxML-NG (v1.2.2) using model GTR+I+G4 with 10 parsimony and 10 random trees. Bootstrapping was performed and reached convergence based on 746 (complete) and 1,000 (reduced) bootstraps. Genetic population relationships were identified using ADMIXTURE (v.1.4.0) with default parameters and K values ranging from 3 to 12.

### Analysis of *HD*, *Mfa*, *STE*, and avirulence gene loci

HD loci from Li et al. (2024) of *Pgt* isolates Pgt21-0 (GCA_008522505.1), 04KEN156/04 (Ug99 GCA_008520325.1), and 99KS76A-1 (PRJNA399035), were used as references. For HD haplotypes, when a long-read assembly was not available, short read assemblies were constructed using Spades 3.10.0 with the -careful option, or minia 3.2.6 with default parameters, and scaffolds were mapped to the known HD loci using Minimap2 2.24 for long-read DNA with the k parameter set to 28 (Li, 2018; Salikhov et al. 2014). This process was repeated with known avirulence gene alleles. Short-read sequences from a total of 371 *Pg* isolates were mapped to a reference fasta file containing all the HD loci. Unique haplotypes with greater than 95% similarity were combined in the classification. Venny was used to display unique haplotypes (Oliveros, 2016). Reads from the same isolates were mapped to genes derived from the Pgt21-0 genome: *Mfa1*, *Mfa2*, *Mfa3, STE3.2–2*, and *STE3.2–3*, with the same methods as the phylogenetic analysis (Luo et al., 2024). Amino acid alignment of the *bW-HD1* and *bE-HD2* from *Pga*, *Pgs*, and *Pgt* were aligned using Geneious alignment with default parameters (v2023.1.2, Biomatters, Auckland, New Zealand). Chimeric haplotypes were analyzed by aligning all haplotypes to individual haplotypes with minimap2 and visualized in Geneious. Network analysis was accomplished using NetworkX (Hagberg et al, 2008) on a set of 86 HD haplotypes aligned using MAFFT 1.5.0 (Biomatters, Auckland, New Zealand). K80 (Kimura 2-Parameter Model), was used to determine the weighted mutation rate. Based on the distance matrix, a threshold of <20% mutation rate differences was chosen for haplotype linkage. Ug99 HD haplotype Pg-bWbE003 was aligned with the *Pgs* haplotypes Pg-bWbE092 and PgbWbE075 separately using Mauve 1.1.3 (Darling et al. 2004). The K-mer Analysis Toolkit (KAT) sect (SEquence Coverage estimator Tool) was used to analyze 99 HD haplotypes from *Pgt*, *Pgs*, and *Pga* (https://github.com/EarlhamInst/KAT; Mapleson et al. 2016). KAT sect was also used to analyze cloned *Pg* avirulence genes. Compatible (*avr*) and incompatible (*Avr*) avirulence gene alleles were postulated based on phenotype data or molecular interactions described in published literature (Arndell et al. 2024; Chen et al. 2025; Lubega et al. 2024; Spanner et al. 2026; Ortiz et al. 2022; Salcedo et al. 2017; Zhao et al. 2022). Avr gene alleles were aligned with MAFFT 1.5.0 (Biomatters, Auckland, New Zealand), and a neighbor-joining tree, generated using the Jukes-Cantor model, was resampled 1000 times to generate bootstrap values.

## Results

### Phylogenetic analysis of *Pg* isolates

Our initial goal was to understand *P. graminis* diversity based on an expanded set of sequenced asexual, sexual, and host adapted lineages. Within the full phylogenetic analysis of 345 *Pg* isolates, *Pga* formed a monophyletic clade distinct from all *Pgt* and *Pgs* isolates (Fig. 1). *Pgs* isolates formed a polyphyletic clade within the larger *Pg* clade, and *Pgt* isolates from a sexual population in Spain formed a clade within the *Pgs* clade. In a reduced phylogeny with a selection of representative diverse isolates, ADMIXTURE analysis at K=3 produced 4 generally distinct populations: a *Pga* only population (orange Fig. S1); a population consisting of mostly Asian, Middle-Eastern, and African isolates, and some North American isolates such as TTTTF (clade 7) from Brazil and USA (light blue Fig. S1); a hybrid Asian/African/North American cluster that included the Ug99 clade 1, one of Ug99’s nuclear parents from clade 2, UN1 isolates from North America, and clade 6-A consisting of TPMKC (74MN1049) and 99EGY6A (light blue and dark blue Fig. S1); the final cluster consisted mostly of North America *Pgt* and *Pgs* isolates, isolates from barley and Mahonia from the Pacific Northwest (PNW), and all four German *Pgs* isolates (dark blue Fig. S1). The Spanish isolates shared a cluster with *Pgs* isolates until K=12. The German *Pgs* isolates consistently shared a minority of their genetics with clade 9 and clade 1 until K=15. The principal component analysis (PCA) of *Pgt* and *Pgs* isolates showed that the variance within *Pgt* isolates is greater than the variance between *Pgt* and *Pgs* isolates (Fig. 2). The full phylogenetic analysis (Figs 1, S2), reduced PCA (Fig. S3), and full PCA (Fig. 2) all suggest that the PNW isolates from barley, such as Epu27, formed their own distinct clade apart from all other isolates. Clade 6A, 7, 6B, and U1 were the only clades to contain North American isolates that fell outside the larger North American (NA) super-clade which contained QFCSC (95MN1080). *Pgt* isolates 87KEN3015D-1, 83ETH13-1, 86MAR179B, 87MAR179B, and 83SYR2-2, were the only non-NA isolates in the NA superclade. The vast majority of Asian, African, and European *Pgt* isolates formed a clade distant from the NA super clade. The NA super clade only contains 18 out of the 99 HD haplotypes described, indicating that NA isolates represent a small fraction of the worldwide stem rust diversity despite being overrepresented in sequencing studies (Figs 3a, 3b; Dataset S2). The phylogenetic analysis strongly supports previously well-established hypotheses that most *Pgt* diversity originates from Asia, and the North American population is heavily bottlenecked.

**Figure 1.**
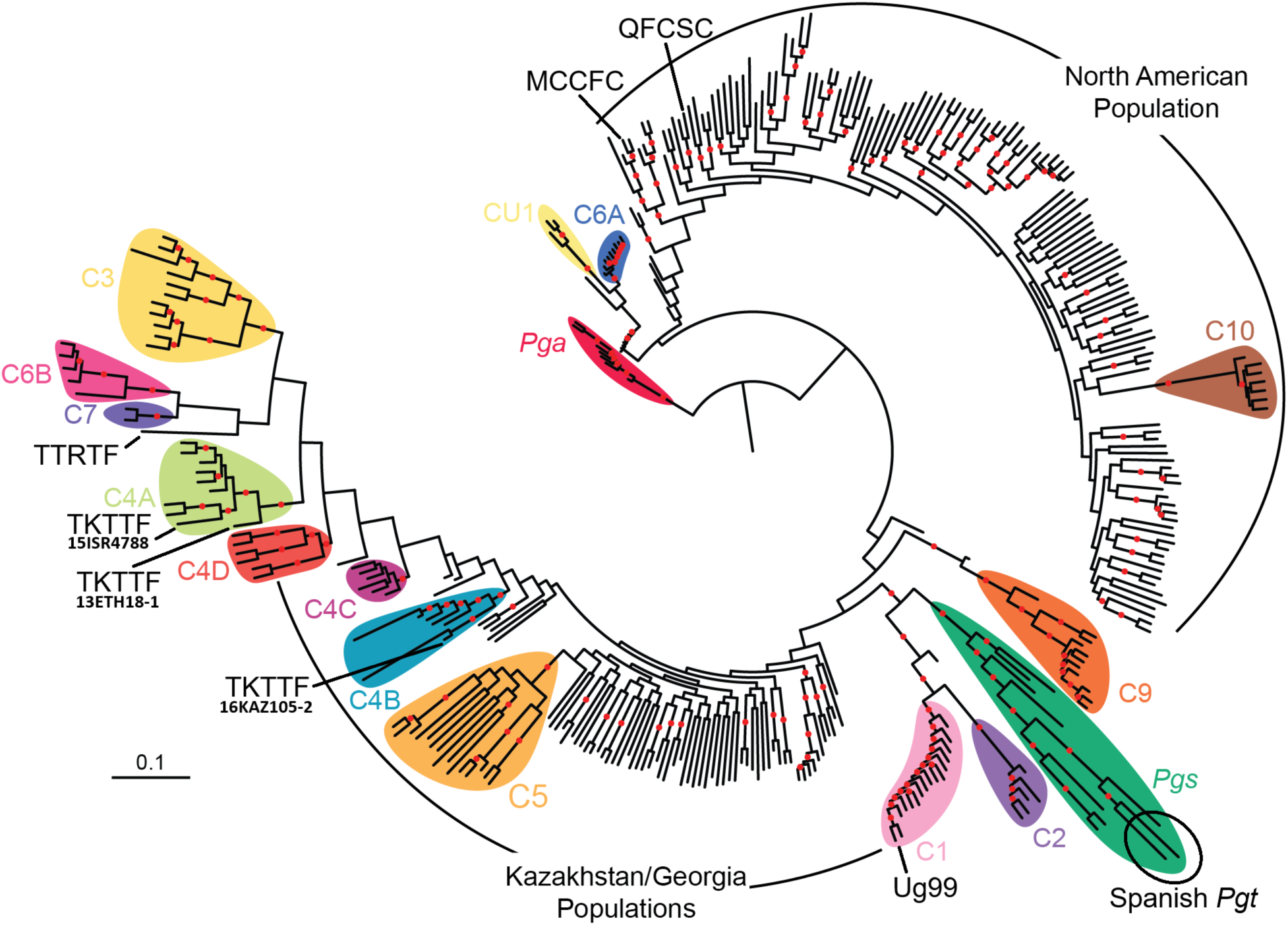
Phylogenetic relationships of major groups of stem rust (*Puccinia graminis*). Read alignment was performed on the *Pgt* isolate 99KS76A reference genome, the initial filtered number of SNPs was 6,585,081 and was reduced to 439,775 based on LD pruning with PLINK2. Known clades are shown with colored labels. Although lacking bootstrap support, the North American and Kazakhstan/Georgian sexual populations are two dispersed genetic groups. Clade 1 contains the Ug99 lineage of wheat stem rust and the sister clade to clades 1 and 2 is rye stem rust (*Pgs*; *Puccinia graminis* f. sp. *secalis*), indicating a close relationship between *Pgs* and Clades 1 and 2. A total of 345 isolates of *P. graminis* were included in the analysis. Oat stem rust (*Pga*; *Puccinia graminis* f. sp. *avenae*) is used as an outgroup. Bootstrap support above 80% is shown as red dots at the center point of branches (based on 746 bootstraps with convergence). Isolate labels for each branch are available in figure S2.

**Figure 2.**
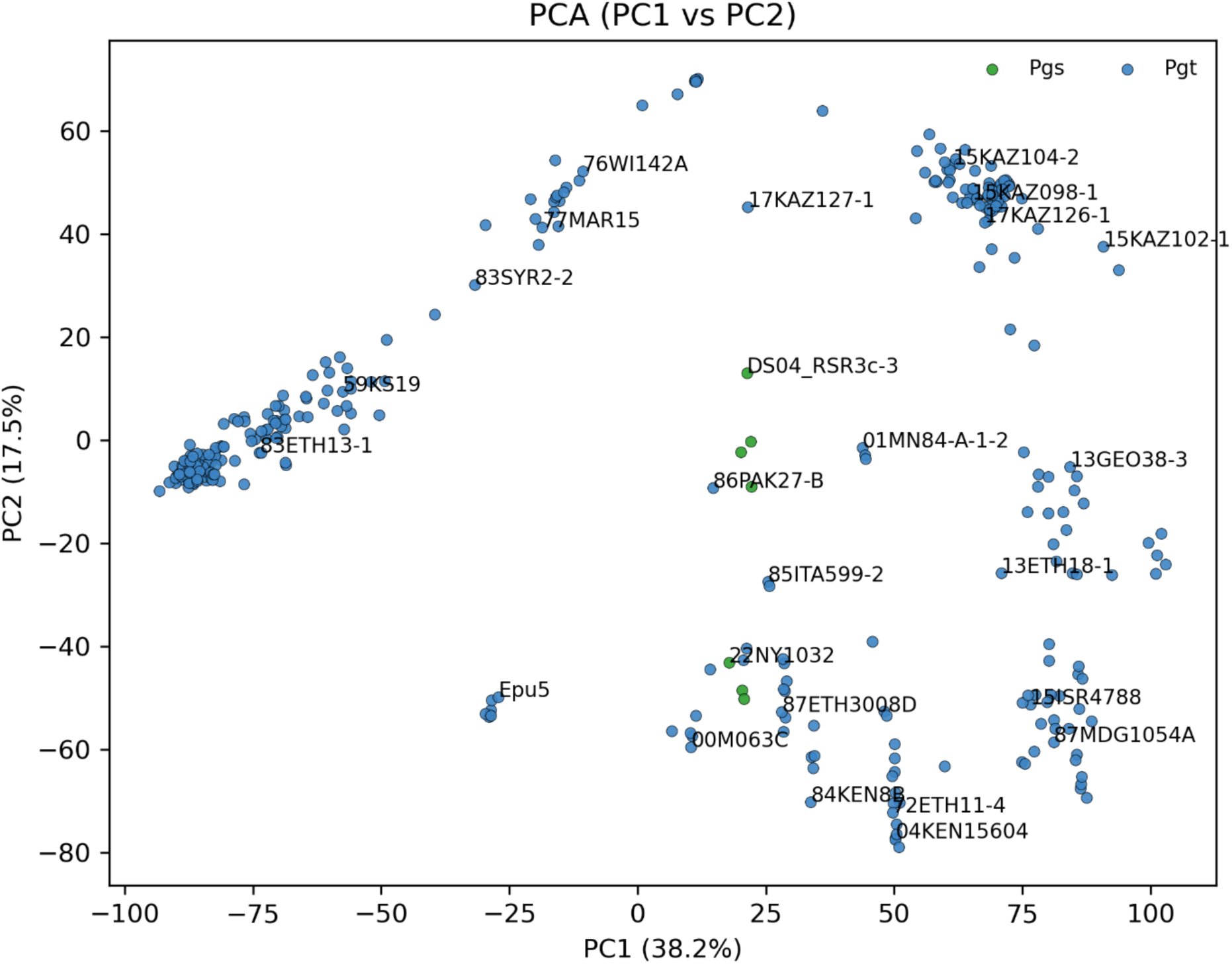
Principal component analysis based on whole genome sequencing. Based on 6,585,081 variants after filtering from *Pgs* (green) and *Pgt* (blue) isolates mapped to the 99KS76A-1 reference genome. Identifiers for a subset of isolates are shown. Host preference determination listed in dataset S1.

**Figure 3.**
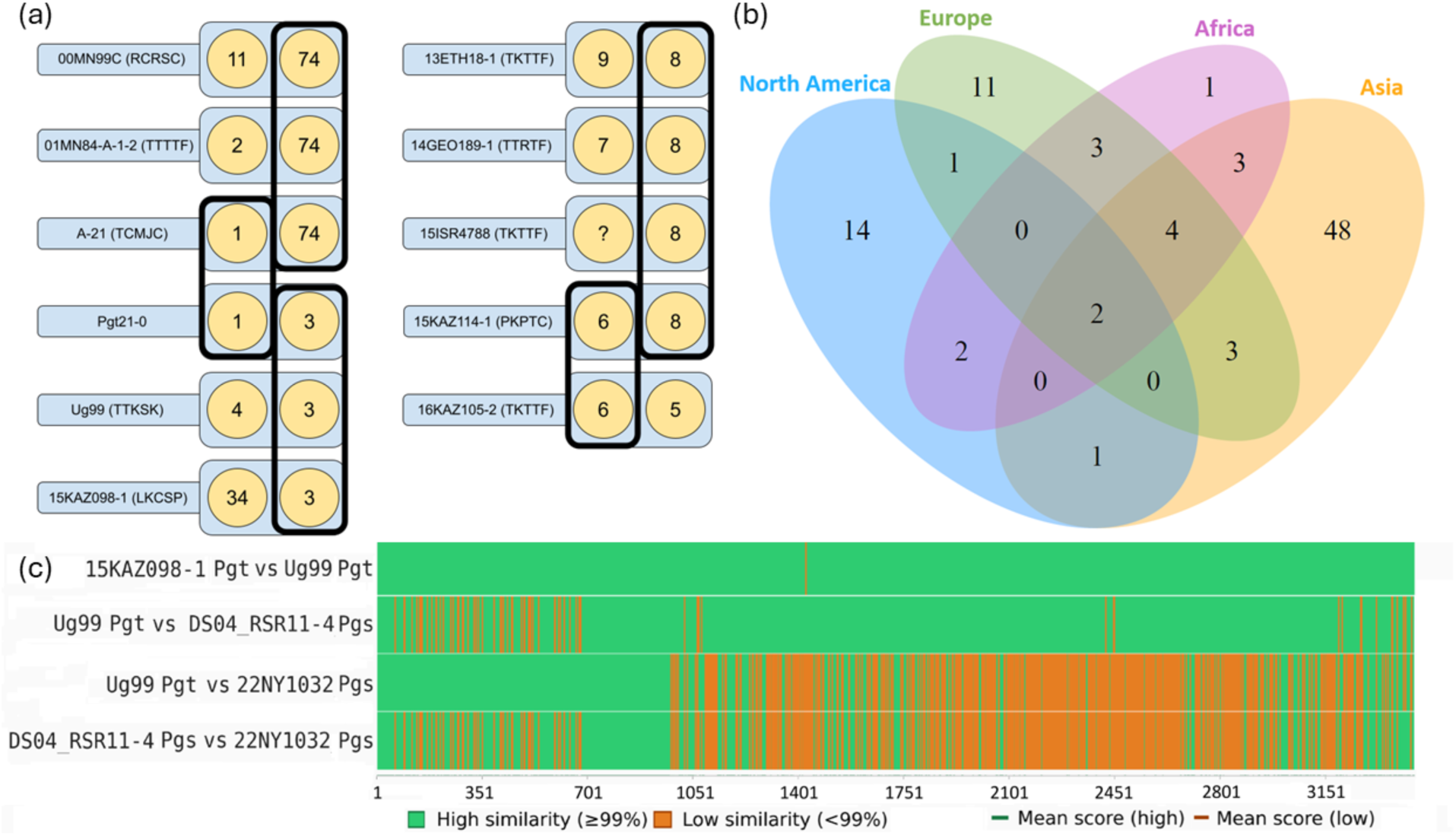
Shared HD haplotypes identify genetic relationships within and between *Pg* lineages. (a) Significant *Pgt* isolates with shared HD haplotypes. Isolate 15ISR4788 (TKTTF) is homozygous for HD Pg-bWbE008. (b) Incidence of HD haplotypes in sequences of isolates collected from four continents determined by >90% containment scores on individual HD haplotypes. (c) Pairwise comparisons of the DNA HD haplotype Pg-bWbE003 of Ug99 (Haplotype A, TTKSK *Pgt*), Pg-bWbE003 15KAZ098-1 (LKCSP *Pgt*), Pg-bWbE092 DS03_RSR11-4, and Pg-bWbE075 22NY1032 (*Pgs*).

### Nuclear exchange in Kazakhstan

Most Kazakh isolates were not present in any previously described clades. *Pgt* isolate 17KAZ127-1 has over 98% containment with the B haplotype of the phased Pgt21-0 reference genome and shares an HD locus with 17KAZ126-1, suggesting it is the result of nuclear exchange, hypothetically through somatic hybridization between a clade 9 isolate and a Kazakh isolate (Dataset S3). Within the larger Afro-Eurasian clade were NA *Pgt* races, TTTTF (01MN84-A-1-2), RCRSC (00MN99C), TCMJC (A-21), and NA *Pgs* isolates. Nuclear exchange is common in *Pgt* isolates from tropical and sub-tropical regions where barberry is less common (Guo et al. 2022). This incidence of nuclear exchange adds this region to a growing list where nuclear exchange in *Pg* has been reported (Guo et al. 2022; Spanner et al. 2026).

### Analysis of *Pg* HD locus haplotypes

The highly divergent basidiomycetes HD mating type loci provide a complementary approach to understand gene flow between *Pg* populations. Due to diversifying selection pressure on the HD locus in heterothallic basidiomycetes, this location contains a large number of highly informative SNPs. Due to *Pgs* and *Pgt* sharing a number of HD haplotypes, the *Pgt*/*Pgs* HD haplotypes are designated with “*Pg*”, while *Pga* haplotypes are designated with “*Pga*”. Highly virulent races frequently share HD haplotypes (Fig. 3). Within *Pg* isolates, the pairwise similarity between HD haplotypes ranged between 61 and 100%, and averaged 70.28% (Dataset S4). The number of isolates sampled from different continents varied greatly and some locations were disproportionately sampled: North America n=151; Africa n=41; Asia n=101; Europe n=18. In total, 99 HD haplotypes were discovered, and a total of 148 HD haplotype combinations. The Pg-bW001-HD1-bE001-HD2 (a 5 kb locus containing *bW-HD1* and *bE-HD2* genes, abbreviated Pg-bWbE001) haplotype is the most common occurring in 21.7% of isolates (n=385) (Dataset S2). In the United States, haplotype Pg-bWbE001 was detected in the earliest NA isolate in the collection, *Pgt* isolate 59KS19 (MCCFC), a representative isolate of Race 56 (Dataset S2). The Pg-bWbE001 haplotype also appeared in Kazakhstan but only in a single *Pgt* isolate 17KAZ127-1 (race CCRSC), which was discovered in southern Kazakhstan near the Tianshan mountains. Both HD loci from *Pgt* isolate 17KAZ127-1 can also be found in isolates in the UN2 clade, and *Pgt* isolate 86PAK27-B (race CCTSC). The second most prevalent HD haplotype was Pg-bWbE103 (11.8%), which was only found in North America. At 11.5% prevalence, three HD haplotypes were found in equal frequency: Pg-bWbE098, Pg-bWbE096, and Pg-bWbE008. In some instances, highly related haplotypes such as PgtF1-107-1-1 (PgtF1) haplotype 11 and 09CA115-2 haplotype 1 were grouped into Pg-bWbE098, as their similarity was >96%. Pg-bWbE098 is also found in an artificially created sexual population from a cross between isolates *Pgt* isolates Pgt4a and Pgt7a (Zambino et al. 2000). Haplotypes similar to Pg-bWbE096 are found in both *Pgs* and *Pgt* isolates (Fig. 3). Pg-bWbE008 was detected in isolates from Germany, Ethiopia, and Kazakhstan. The Pg-bWbE074 allele was only detected in race TTTTF (found in Brazilian isolate 81BRA38A and US isolate 01MN84-A-1-2), clade VI-A, and isolate A-21 (TCMJC, a race notable for defeating *Sr13*) from North Dakota (Upadhaya, Upadhaya, & Brueggeman, 2024) (Fig. 3a). The three Spanish isolates formed their own clade with *Pgs* and carried distinct HD haplotypes with no direct similarity to other HD loci. Due to limitations of short read *de novo* assemblies, not all HD haplotypes extracted from these assemblies span both *bW-HD1* and *bE-HD2* (Table S1). The large diversity of HD haplotypes in Asia supports the previous conclusion that Asia contains a greater diversity of *Pgt* isolates than other continents, and North America possibly has one of the least diverse populations (Bradshaw et al. 2024).

### Diversity of mating genes

Previous studies have indicated that the genes in the HD locus are under a much greater diversifying selection pressure due to the need for HD haplotypes to be different to form a functioning dikaryotic cell. Amino acid alignment showed regions of variability and stability within the *bW-HD1* and *bE-HD2* genes located in the HD haplotype region (Fig. S4). Within 305 *Pga*, *Pgs*, and *Pgt* isolates mapped to *Mfa1* (204 bp), *Mfa2* (102 bp), *Mfa3* (171 bp), *STE3.2-2* (1167 bp), *STE3.2-3* (1140 bp), there were a total of 18, 5, 24, 72, and 75 single nucleotide polymorphisms (SNPs) respectively (Table 1, Fig. S5-S9). Matrices of 66 aligned *bW-HD1* and *bE-HD2* nucleotide coding sequences, with the lowest pairwise similarities between genes being 65.34% and 73.75% respectively, and the average similarity for all 66 haplotypes being 70.28% (Dataset S4). The diversifying selection pressure (dN/dS non-synonymous/synonymous mutation ratio) is 2.92 for *bW-HD1* (n=67) and 2.82 for *bE-HD2* (n=83) (Table 1). These mutation ratios clearly indicate the HD locus is mutating at a much higher rate than the pheromone precursor and pheromone receptor genes.

**Table 1.** Single nucleotide polymorphisms present in the coding DNA sequences of mating type genes. Pheromone precursor (*Mfa*) pheromone receptor (*STE*) and homeodomain transcription factor genes (*HD*) in *Pga*, *Pgs*, and *Pgt* haplotypes extracted from short-read *de novo* assemblies. Non-synonymous/synonymous mutation ratio (dN/dS) reported in *Pgs+Pgt* or *Pga+Pgs+Pgt* (All *Pg*) haplotypes.

| Gene | *Mfa1* | *Mfa2* | *Mfa3* | *STE3.2-2* | *STE3.2-3* | *bW-HD1* | *bE-HD2* |
| --- | --- | --- | --- | --- | --- | --- | --- |
| CDS length (nt) | 204 | 102 | 171 | 1167 | 1140 | 1851 | 1170 |
| *Pgs+Pgt* dN/dS | 1.2 | -* | 3 | 0.56 | 1.9 | 3 | 2.8 |
| *Pgs+Pgt* SNPs nt | 11 | 1 | 4 | 11 | 143 | 1113 | 694 |
| *Pgs+Pgt* SNPs % | 5 | 0.98 | 2.3 | 0.94 | 12.5 | 60.1 | 59.3 |
| All *Pg* dN/dS | 0.78 | 1 | 2.6 | 0.3 | 1.2 | 2.9 | 2.8 |
| All *Pg* SNPs (nt) | 25 | 6 | 29 | 78 | 204 | 1349 | 699 |
| All *Pg* SNPs (%) | 12.2 | 5.8 | 14 | 6.1 | 17.9 | 72.8 | 58.7 |

\*No non-synonymous mutations detected.

### Chimeric HD haplotype in UgGG links *Pgs* and *Pgt* lineages

To investigate potential evidence of historical sexual recombination in the HD locus, an analysis was performed on chimerism among HD alleles. Two chimeric haplotypes aligned to the Ug99 haplotype Pg-bWbE003. Pg-bWbE075, a haplotype uniquely found in *Pgs* isolates which was derived from the *Pgs* isolate 22NY1032 shares 1,063 bp at 100% identity with Pg-bWbE003, The Pg-bWbE092 haplotype from DS04_RSR11-4 shares a 2,746 bp region with 99.6% identity to Ug99 Pg-bWbE003 haplotype, with several missing regions in the alignment (Fig. 3c). This suggests that one haplotype of the HD locus in Ug99, or a progenitor of Ug99 or Clade 9, emerged through a hybridization event between *Pgt* and *Pgs*.

### Admixture and ancestry analysis of *Pgs* and *Pgt* HD haplotypes and avirulence genes

The role of ancestral sexuality between *Pgt* and *Pgs* prompted an examination of evidence within whole genome sequencing data of admixture between these host adapted lineages. Network analysis revealed two clusters that differentiated at a threshold of 0.2. *Pgt*, *Pgs*, and *Pga* were all present in a large cluster containing 83 HD haplotypes (Fig. 4). *Pga* haplotypes 1, 2, and 3 only connected to a single *Pg* haplotype Pg-bWbE015. Pga-bWbE004 shared no close ancestry with the three other *Pga* haplotypes but did have ancestry with *Pgt* haplotypes Pg-bWbE001 and Pg-bWbE065. *Pgt* HD haplotypes Pg-bWbE004, Pg-bWbE044, and Pg-bWbE050 formed their own cluster distinct from the larger cluster (Fig. 4).

**Figure 4.**
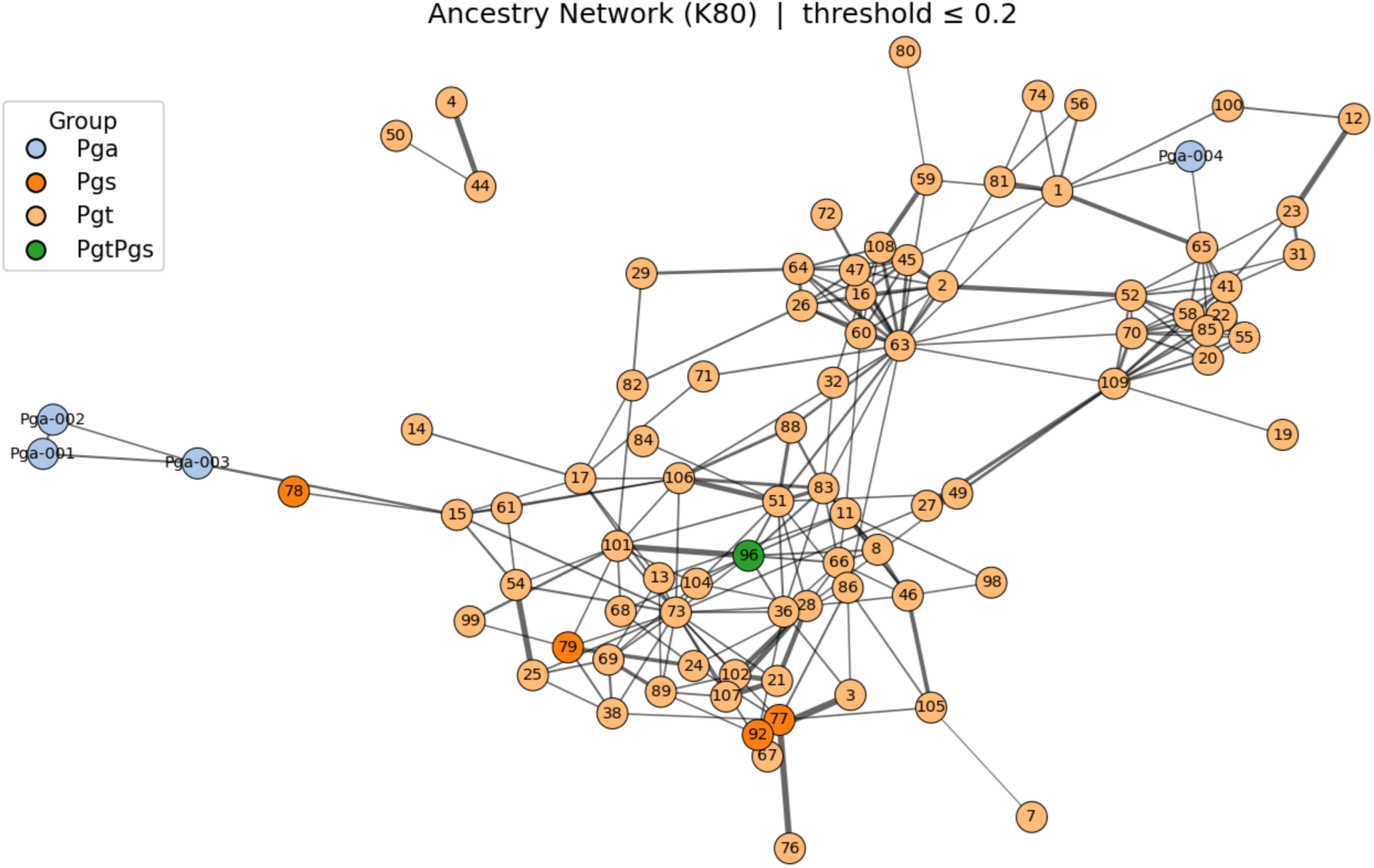
Network analysis of unique *Puccinia graminis* HD haplotypes from wheat, rye, and oat stem rust. Total number of haplotypes in analysis is 86. Haplotype names for *Pg* HD haplotypes abbreviated to the haplotype number. A weighted model, K80 (Kimura 2-Parameter Model), was used to determine mutation rate. Haplotypes with >95% similarity were removed. Network links indicate <20% DNA mutations per site differences between haplotypes.

K-mer analysis of known *avr* alleles showed that in addition to sharing HD haplotypes, there was gene flow between *Pgs* and *Pgt* around the world and in almost every *Pgt* clade (DatasetS2, S5). *Pgs* isolates contain known compatible (virulent) alleles of *avrsr13a*, *avrsr13d*, *avrsr27*, *avrsr33*, *avrsr50*, and *avrsr62*. Phylogenetic and containment analysis of avirulence gene alleles strongly suggest compatible alleles derived from *Pgs* of *avrsr13a*, *avrsr13d*, *avrsr22*, *avrsr33*, and *avrsr50* are responsible for *Pgt* isolates gaining virulence to wheat lines containing the corresponding resistance genes (Figs 5, S10-17; Dataset S5). For example, the *Pgt* isolate QCMJC (07WA140) causes disease on plants containing *Sr50*, and the allele of *avrsr50* derived from that isolate is directly derived from *Pgs* (Fig. 5) (Chen et al. 2025; Spanner et al. 2026; Ortiz et al. 2022).

**Figure 5.**
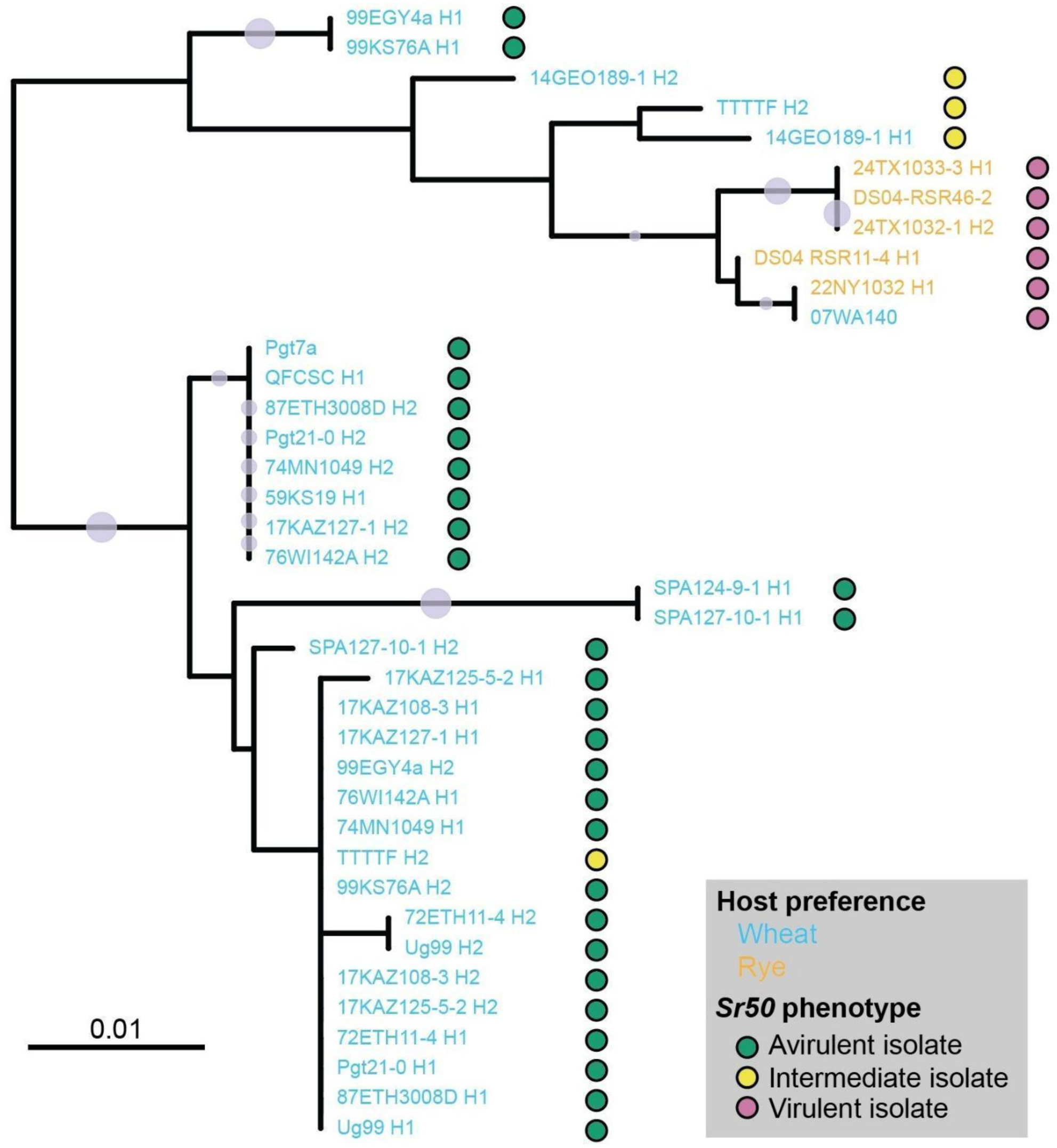
Phylogenetic tree of *AvrSr50* alleles derived from *Pgs* or *Pgt* isolates and their ability to cause disease on *Sr50*. Alleles of AvrSr50 derived from *Pgs* and *Pgt* are denoted in orange and blue, respectively. Avirulent = fleck to 2, intermediate = 2+ to 3-, virulent = 3 to 4 disease phenotypes. Tree was generated using alleles that were identified in short read *de novo* assemblies or long-read reference genomes. Alleles were aligned with MAFFT, and a neighbor-joining tree, generated using the Jukes-Cantor model, was resampled 1000 times to generate bootstrap values.

## Discussion

### *Pg* admixture, gene flow, and predictions on the breakdown of resistance genes in wheat

Currently, detecting virulence on cultivated plant hosts requires retrospective studies on outbreaks and epidemics. With genomics, it is becoming increasingly possible to determine the stability of resistance genes by sequencing pathogen populations and understanding the underlying mechanisms of gain of virulence events. In this study we attempted to illuminate the gene flow of virulence genes from non-wheat adapted *Pg* strains into *Pgt*. Previous studies on specialized forms of stem rust revealed that F1 offspring of crosses between *Puccinia graminis* from wheat, oat, rye, and bentgrass yielded isolates with larger host ranges, but overall less virulence and infrequent fertility (Johnson, 1949). In this work, several lines of evidence support the role of gene flow between *Pgt* and *Pgs* populations as a basis for the emergence of virulent lineages of stem rust. First, *Pgs* and *Pgt* were readily distinguishable from *Pga*, while *Pgs* and *Pgt* isolates could not be consistently separated phylogenetically (Figs 1, 2). *Pgs* is located within a large *Pgt* clade that includes mostly Asian, African, and European isolates, and the Spanish *Pgt* isolates are located within the *Pgs* clade. Second, mating genes and cloned secreted effectors are shared between *Pgs* and *Pgt,* but neither share these genes with *Pga*. Importantly, the emergence of virulent lineages of *Pgt* are associated with alleles of effectors recognized by resistance genes likely derived from *Pgs*. For example all *Pgt* isolates virulent on *Sr50* contain alleles of *avrsr50* that are otherwise exclusive to *Pgs* (Fig. 5). *Pgs* isolates, such as 22NY1032, 24TX1033, and 24TX1032 are unable to form pustules on any of the wheat lines in the current wheat stem rust differential set, which contains 24 monogenic lines in various wheat backgrounds, but these *Pgs* isolates will infect “Winter Rye” and “Prolific” rye lines in a panel of standardized lines of highly susceptible wheat, barley, oat, rye, and wild grasses to capture host preference of rust isolates. Virulence on *Pgt Sr* genes is essentially “hiding” inside non-wheat adapted stem rust such as *Pgs*. The presence of virulence alleles for almost every cloned avirulence gene indicates that not only is this phenomenon possible, but is actively causing gain-of-virulence events. Both wheat and grass resistance genes found in wheat for stem rust are at risk. Breakdowns in resistance often seem to come out of nowhere, but in reality, virulence is simply waiting for the correct selection pressures to emerge. The recent Kazakh and Spanish *Pgt* races are important examples of this phenomenon, being populations that have emerged novel virulences on important previously stable resistance genes.

### *Pg* diversity in Asia and the formation of recent epidemics

The threat of stem rust to cereal production lies in its ability to cause complete yield loss under severe infection. Sexual recombination and nuclear exchange are the primary methods by which novel virulence combinations (races) are produced in the genus *Puccinia* (Hovmøller et al., 2026; Guo et al., 2022). This study was initially motivated by the goal to understand the origin and global context of the Kazakhstan outbreak that started in 2015 and the role of gene flow and admixture events globally. The outbreak was caused by a phylogenetically distinct, highly diverse sexual population of *P. graminis* f. sp. *tritici* (Olivera et al., 2022; Shamanin et al., 2020). This population highlights the importance of controlling *Berberis* spp. to limit the production of novel races in areas near wheat production. With the movement of Ug99 into Iraq (Nazari et al. 2021) and Nepal (Patpour et al., 2024), there is a significant risk of Ug99 and other highly virulent lineages recombining with sexual strains of *P. graminis* (*Pg*) in Asia and Europe to create novel and hypervirulent races that could undermine wheat production. Over 50% of the world’s wheat production occurs in countries near Central-Asia, and wind-borne pathogens can travel thousands of kilometers in a single season (FAO, 2022).

It is widely thought that *Pg* is native to central Asia, where both its hosts, barberry and susceptible grass species, are native and abundant (Bradshaw et al., 2024). Asia contains the highest number of HD haplotypes, although this result is based on unequal sampling. Both the central Asian and Spanish populations represent previously undescribed reservoirs of cryptic ancient lineages of *Pg* due to the presence of HD haplotypes unique to these regions. Previous studies on *P. striiformis* f. sp. *tritici* (*Pst*) have shown the efficacy of using the HD locus as an initial benchmark to understand epidemics (Holden et al., 2023). As a genetic marker, the HD locus is a permanently heterozygous, highly divergent locus that can be used to understand pathogen evolution at a highly specific level. The HD locus provides researchers with the opportunity to examine molecular footprints of *Pg* populations and understand the impact and depth of the introduction of novel strains into native populations.

A significant challenge in understanding the origin, hybridization, and movement of cereal rusts is a lack of long-term temporal, comprehensive worldwide pathogen sampling. Comprehensive sampling of the Georgian and Kazakh/Siberian populations identified several striking patterns and connections to other epidemics. Significantly, we have demonstrated a significant amount of gene flow between Africa and Central Asia, especially regarding races that have caused epidemics such as Ug99 and TKTTF. The HD locus provides important evidence in understanding the level of genetic variation within *Pg* and gene flow between *Pg* populations. The Georgian sexual population shared HD alleles with the ubiquitous HD alleles found in Clade 4 and Kazakhstan. Thus, the Asian sexual populations may be the origin of the Clade 3 and 4 races that caused epidemics in Ethiopia (Olivera et al., 2015) and outbreaks in Georgia (Olivera et al., 2019) and Germany (Olivera Firpo et al., 2017). Using spore dispersal models, Meyer et al., (2017) also suggest that races originating in the Caucasus region and West Asia can spread into East Africa and Europe. German isolates from 2013 had HD alleles exclusively found in Georgia and Kazakhstan, although it is difficult to determine directionality of genetic migration. *Pgt* isolate 13ETH18-1 (race TKTTF) is found in clade 4A and caused the severe epidemic in Ethiopia on cultivar ‘Digalu’ (carrying *SrTmp*) (Olivera et al., 2015). This isolate shares HD haplotypes with Georgian and German isolates and shares the HD haplotype Pg-bWbE008 with four Kazakhstan isolates from 2015, two 2016 isolates, two 2017 isolates, and the clade 3B race TTRTF (Dataset S2). However, *Pgt* race TKTTF from Kazakhstan (16KAZ105-2) is phylogenetically distinct from the Ethiopian isolate, and both HD alleles are different. The second HD allele of *Pgt* race TTRTF is unique to that isolate, indicating this race was formed by recombination or a somatic hybridization event with an isolate with the ‘Digalu’ Pg-bWbE008 haplotype, and an unknown *Pg* isolate. All but one of these isolates containing Pg-bWbE008 is virulent on *Sr3C*. This HD haplotype was present in an Israeli isolate from 2012 (12ISR2083), which is the earliest sequenced isolate containing this HD haplotype, although isolates with similar phenotypes have existed in the Middle East since 2005. Sequencing of available historical isolates would be needed to make a conclusion about the origins of the ‘Digalu’ outbreak, but certainly Ethiopian, Georgian, German, and Kazakh populations are all genetically connected.

Notably, several TKTTF isolates have only one HD haplotype and appear to be homozygous for Pg-bWbE008 (Dataset S2), introducing the possibility that an isolate could exchange nuclei with itself via somatic hybridization, or the production of trinucleate urediniospores which could lead to a homokaryotic *Pg* isolate (Nelson et al. 1955; Nelson, 1956). This further raises the possibility that a single heterokaryotic dikaryon could form two more distinct races by producing two homokaryotic *Pg* isolates, each one having two copies of one of the nuclei present in the heterokaryotic individual. A single isolate could, hypothetically, evolve into a total of three race phenotypes in a single growing season. The rate at which somatic hybridization occurs is not well studied, although in Africa it appears to be the most likely means of genetic exchange between isolates. Given that not every HD locus is likely self-compatible, and homokaryotic *Pg* isolates may have reduced fitness, it is probable that these individuals are relatively rare.

Our ability to understand and predict disease epidemiology requires comprehensive knowledge of the genetics and virulence of a pathogen. The isolates derived from the stem rust samples collected in Kazakhstan yielded substantially high numbers of unique HD haplotypes, likely due to considerable genetic diversity and the yearly formation of novel recombinant strains. Given the widespread prevalence of the alternate host species, admixture in Kazakhstan and Siberia will likely continue to contribute to stem rust epidemics in the future. Both new races, RCMTM (17KAZ125-4-2) (virulent on *Sr47*) and RHHTM (17KAZ108-3 virulent on *Sr32* and *Sr40*), are from previously unsequenced genetic backgrounds with novel HD loci only found in Kazakhstan. The observation that several recently sequenced isolates with novel race types contain novel HD haplotypes underscores the need for more global monitoring and sequencing of *Pg* isolates particularly from natural sexual populations.

### Rye and wheat stem rust admixture linked to *Sr31*, *Sr22, Sr27*, and *Sr50* virulence

Several stem rust resistance genes of rye origin, including *Sr27*, *Sr31*, *Sr50*, *Sr59*, and *Sr1RS_Amigo*, have been introgressed into wheat lines providing strong and stable resistance against *Pgt* (Roelfs 1985). The popularity of the wheat/rye hybrid Triticale has also risen over the last century, from 250,000 acres in 1960 to over a million acres today in the USA (Oelke et al. 1989). This study has outlined multiple instances of admixture events between the wheat and rye stem rust pathogens that are directly linked to a breakdown in rye-derived resistance genes. Similar breakdowns in resistance, due to breeding efforts increasing interspecies hybridization, have been observed in rye and wheat powdery mildew (Menardo et al. 2016). *Sr50* virulence developed in at least three independent events, and each time, an allele previously exclusive to rye stem rust isolates was underlying virulence. Virulence on *Sr50* appears likely the result of admixture between the wheat and rye stem rust pathogens in sexual populations. The Spanish *Pgt* isolates form a unique clade that appears to have substantial admixture with *Pgs*, which is likely a significant source of diversity for this population. While Ug99 has less phylogenetic and admixture relationship to *Pgs* than the Spanish isolates, the similarity of one of its HD haplotypes to that of two rye stem rust isolates points strongly to a recent sexual event that may have formed Clade 1 or Clade 9 (Fig. 3c). Clade 1 is a sister clade to *Pgs*, but there is no obvious direct ancestor to either group that would explain the true origin.

Introduction of *Pgs* allelic variants of *avr* genes was not restricted to rye-derived resistance genes. *Sr22* is derived from *Triticum monococcum* and the introduction of *Pgs avrsr22* into *Pgt* appears to be coincidental. The closest ortholog to the wheat *Sr13* (*T. monococcum*) in rye is only 35% similar (ALO61078), so again *Pgs* containing compatible *avrsr13a* and *avrs13d* alleles appears to be coincidental. The gene *Sr33* (*Aegilops tauschii*) in wheat has an ortholog in rye (ALO61079) that is 86% similar. Given that this study only examined cloned avirulence genes, it is highly likely that there are many more resistance genes which have been broken by *Pgs*, or other *Pg formae speciales*.

### Sexual populations in North America and *Pgs* hybrids

Barberry eradication laws in the United States have played a key role in reducing the diversity of *Pgt* and severity of stem rust in North America. The first recorded instances of barberry eradication laws date back to 1754 in New England (Davis, 1907). However, sexual populations of *P. graminis* have thrived in the PNW of the United States, mostly due to naturalized barberry and native plants in the Berberidaceae family such as *Mahonia* spp. that host the fungus’ aecial stage on their fruits (Jin et al., 2014). A survey was conducted in the 1970s that resulted in 100 novel races in 426 sexual isolates discovered in the United States (Roelfs & Groth, 1980). Another more recent study collected isolates from the states of Washington, Idaho, and North Dakota, with most of the Washington and Idaho isolates collected in 2019 and 2020 (Upadhaya et al. 2024). The isolates in that study were mainly derived from samples collected from barley (a common host for both *Pgt* and *Pgs*) and *Mahonia*. These recent isolates were sequenced, enabling detection of significant GWAS markers for *AvrSr5*, *AvrSr21*, *AvrSr8a*, *AvrSr17*, *AvrSrSa*, and *AvrSrSd*. According to both the phylogenetic and HD analysis, much of this population is the result of admixture of *Pgt* and *Pgs*. Barley is an excellent host for these types of admixture events to occur, as most resulting hybrids would have lower virulence on both rye and wheat, and would generally be selected against in the short term by rye or wheat hosts. There are 13 Washington state isolates from 1970s, one California isolate from 2009 (09CA115-2, BCCBC), and 26 recent isolates from the Upadhaya et al. (2024) PNW population containing both *Pgt* and *Pgs* (Pg-bWbE096) HD haplotypes. These two forms of *P. graminis* were previously delineated by host preference, with the majority of *Pgs* isolates able to infect rye and barley, but unable to infect wheat, and vice versa. The wheat line McNair 701 is often used as the susceptible check in wheat stem rust phenotyping (Roelfs & McVey, 1979). However, *Pgt* races such as QCCJB (isolate 01SD80A from barley) produced a low infection type on this variety, indicating additional resistance genes in this genetic background that make wheat cv. McNair 701 less than ideal for capturing both *Pgt* and *Pgs*. PI 532013 (barley), *Elymus glaucus* (blue wildrye), and wheat cv. Line E are recognized as universally susceptible to rye and wheat stem rust pathogens (Jin, 2005). While the underlying genetics contributing to the host preference of *Pgt* and *Pgs* is currently unknown, it may be a similar model of high frequency species-wide resistance genes underlying host species specificity observed in barley-*P. striiformis* f. sp. *tritici* interactions (Bettgenhaeuser et al. 2021).

If admixture between *Pgt* and *Pgs* becomes more common, it is likely that novel races could emerge as the resistance genes preventing *Pgs* from infecting wheat are lost. Alternatively, *Pgt* and *Pgs* may be evidence of strong selective pressure exerted by large-scale planting of crops with narrow genetic diversity in contemporary agriculture systems. A significant challenge with understanding virulence in rye stem rust is the lack of a rye stem rust differential set as there is in wheat. Therefore, to date the classifications of stem rust isolates are based on their derived host (i.e. wheat or rye). The work presented here highlights the importance of further scientific research on rye stem rust due to its role in gene flow with *Pgt*. While admixture between *Pgs* and *Pgt* seems somewhat common, especially in the PNW, there is no evidence that *Pga* is recombining with any other *formae speciales* evaluated in this study.

There are other signs that while the dominant race in North America, QFCSC, continues to be present in wheat field year after year, unrelated stem rust strains persist in many regions in North America, and the next outbreak may emerge from these populations. *Pgt* isolate A-21 (TCMJC) from North Dakota (Upadhaya et al. 2024) had Pg-bWbE074, a haplotype only found in *Pgt* race TTTTF or clade 4A isolates (RCRSC). It was previously proposed that *Pgt* race TTTTF was likely the parent of race RCRSC, and the HD analysis supports that hypothesis (Guo et al., 2022). A subset of isolates within the PNW also contain a novel HD haplotype Pg-bWbE100 found in the Epu isolates from barley in Pullman, Washington, USA between 2019-2020. The second HD haplotype in these isolates was *Pgt* HD haplotype Pg-bWbE085 found in isolate 01SD80A (race QCCJB) (Dataset S2), an isolate low on the highly susceptible wheat line McNair 701, but highly virulent on barley (Solanki et al., 2019). While barley adapted stem rust (*Pgh*) is largely unstudied by the cereal rust community, it may be playing an important role as an intermediate host for *Pgt*/*Pgs* hybrids. During an initial cross between *Pgs* and *Pgt* isolates, the F1 population would almost certainly be weakly virulent on rye or wheat. This is compounded by the fact that despite its popularity, wheat is mostly grown in regions with little to no barberry in the USA. The hybridization of *Pgt* isolates with undescribed *Puccinia* lineages represents an additional threat for managing *P. graminis*.

## Conclusion

Cereal rusts, and stem rust in particular, represent one of the greatest threats to calorie production in the world. As novel races spread, sexual recombination will continue to play a role in increasing the genetic diversity of *P. graminis*. Cryptic populations of *P. graminis* are responsible for several of the hypervirulent strains found since the Ug99 outbreak. Given how few *Pgs* isolates have been sequenced, it is also possible that recombination with these adapted lineages plays a larger role in gain of virulence events in *Pgt* than previously recognized. The mapping of HD haplotypes has provided valuable insights into the admixture events in *P. graminis*. While this type of analysis is limited to the small number of sequenced *Pg* isolates, it explains many patterns seen within that past 100 years of researching *P. graminis* isolates. These analyses show a risk that virulence on *Sr13*, *Sr22*, *Sr27*, *Sr33*, and *Sr50* could arise in North American *Pgt* isolates given admixture with *Pgs*. This research highlights the importance of genomics in tracking epidemics, and the need for sufficient sampling and sequencing to help researchers understand host-pathogen interactions.

## Acknowledgements

The authors would like to thank Kim-Phuong Nguyen, Les Szabo, Jerry Johnson, Krista Ristinen, Sam Stoxen, Merlin Van Alstine, Sam Gale, and Thai Lee for their significant contributions to this research. Funding for this project was provided by the Gates Foundation (INV-004430), USDA-ARS (5062-21220-025-000D), USDA Non-Assistance Cooperative Agreement (58-5062-1-014), and CIMMYT. This research used resources provided by the SCINet project and/or the AI Center of Excellence of the USDA Agricultural Research Service, ARS project numbers 0201-88888-003-000D and 0201-88888-002-000D. Computational resources were provided by the USDA SCINet Ceres cluster, and the University of Minnesota-Minnesota Supercomputing Institute (MSI).

## Author Contributions

MR, YJ, PO, EA, AA, MM, and TI conceived and planned the project. AA prepared the libraries, provided QC, and sequenced DNA samples. TI, MR, YJ, and PO single pustuled and phenotyped the isolates. TI, MV, MR, and MM analyzed the data and prepared the publication. All authors contributed to the final draft of the publication.

## Data Availability

All new sequencing data described in this manuscript has been deposited in NCBI BioProject PRJNA1263545. Other sequencing data sets used for analysis include NCBI BioProject PRJEB47693, PRJNA803546, and PRJNA952228. Multiple sequence alignments, phylogenetic trees, and associated scripts are available on Figshare (https://doi.org/10.6084/m9.figshare.34364202). Generative AI were used including ChatGPT v5.6 Sol were used to develop R scripts for data visualization.

## Supporting Information

Figure S1. Phylogenetic and population structure analysis of a representative set of *Pg* isolates.

Figure S2. Phylogenetic relationships of major sexual populations of stem rust (*Puccinia graminis*).

Figure S3. Principal component analysis of the *Pg* isolates using 9,677,946 SNPs of variants called on the Pgt21-0 genome.

Figure S4. Amino acid alignments of (A) *Pg bW-HD1* (B) and *Pg bE-HD2* genes.

Figure S5. Neighbor joining tree developed with the Jukes Cantor model, replicated 1000 times with bootstrap values >80 reported of 100 *Pg* HD haplotypes.

Figure S6. *Mfa1* neighbor-joining tree developed with the Jukes-Cantor model based on 18 DNA SNPs.

Figure S7. *Mfa2* neighbor-joining tree developed with the Jukes-Cantor model based on 5 DNA SNPs.

Figure S8. *Pra STE3.2-2* Pgt21-0 neighbor-joining tree developed with the Jukes-Cantor model based on 72 DNA SNPs.

Figure S9. *Pra STE3.2-3* Pgt21-0 neighbor-joining tree developed with the Jukes-Cantor model, based on 75 DNA SNPs.

Figure S10. Phylogenetic tree of *AvrSr13* alleles and postulated compatibility on *Sr13*.

Figure S11. Phylogenetic tree of *AvrSr22* alleles and postulated compatibility on *Sr22*.

Figure S12. Phylogenetic tree of *AvrSr27* alleles and postulated compatibility on *Sr27*.

Figure S13. Phylogenetic tree of *AvrSr33* alleles and postulated compatibility on *Sr33*.

Figure S14. Phylogenetic tree of *AvrSr35* alleles and postulated compatibility on *Sr35*.

Figure S15. Phylogenetic tree of *AvrSr50* alleles and postulated compatibility on *Sr50*.

Figure S16. Phylogenetic tree of *AvrSr62* alleles and postulated compatibility on *SrC2*.

Table S1. HD haplotypes contributing to novel *Pg* races, clades, or epidemics.

Dataset S1. Sequencing inventory and phenotypes of isolates included in this study.

Dataset S2. Kat sect containment of sequenced isolates on HD haplotypes.

Dataset S3. Containment of Kazakh isolates 17KAZ127-1 and 15KAZ098-1 against long read assemblies of Pgt21-0 and Ug99 respectively.

Dataset S4. Distance matrix of aligned *Pg* HD haplotypes.

Dataset S5. Containment of all sequenced isolates on postulated/cloned Avr genes.

