## Supporting Information for "Rye stem rust is a reservoir of diversity and virulence for wheat stem rust"

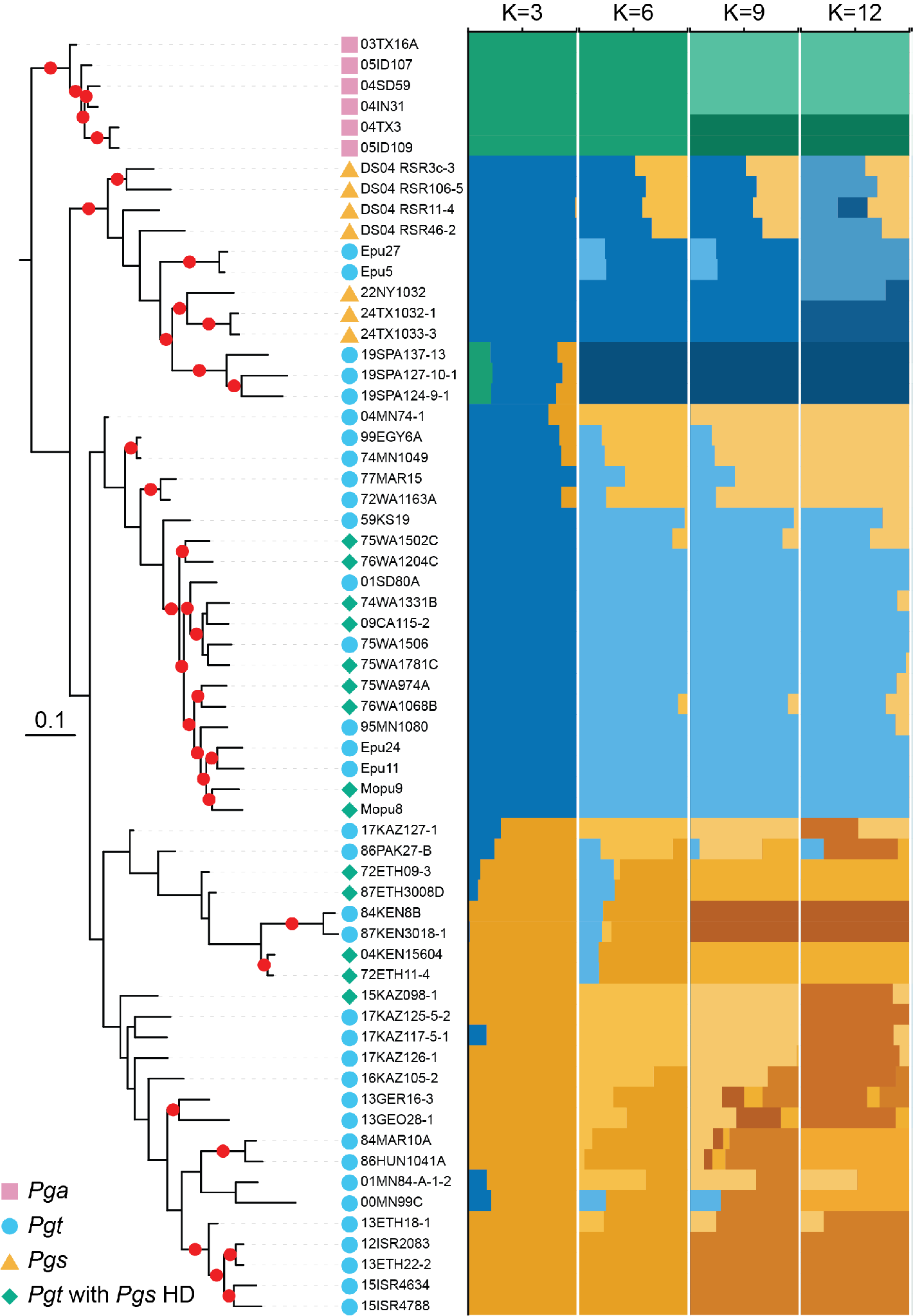

**Figure S1. Phylogenetic and population structure analysis of a representative set of *Pg* isolates.** Read alignment was performed on the *Pgt* isolate 99KS76A reference genome, the initial filtered number of SNPs was 6,585,081 and was reduced to 633,816 based on LD pruning with PLINK2. A total of 62 isolates of *P. graminis* were included in the analysis. Oat stem rust (*Pga*; *Puccinia graminis* f. sp. *avenae*) is used as an outgroup. Host status is denoted by colored shapes in the key. Bootstrap support above 80% is shown as red dots at the center point of branches based on 1,000 bootstraps with convergence. Right panel, ADMIXTURE results based on K values of 3, 6, 9, and 12. Color coding denotes maximum likelihood estimates of individual ancestries based on the same SNPs used for phylogenetic tree construction.

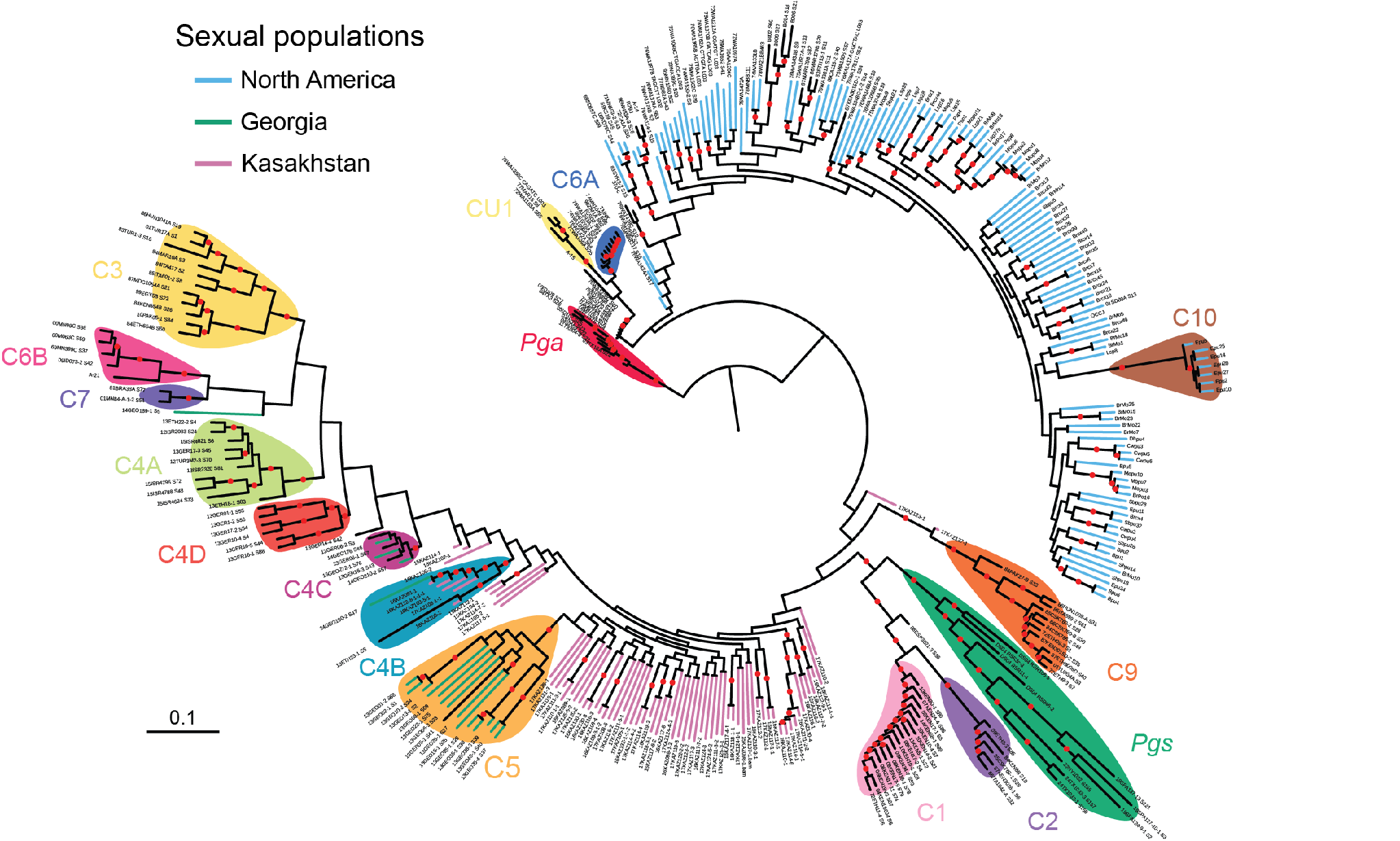

**Figure S2. Phylogenetic relationships of major sexual populations of stem rust (*Puccinia graminis*).** Fig. 1 overlaid with sexual population status and isolate identifiers. Read alignment was performed on the *Pgt* isolate 99KS76A reference genome, the initial filtered number of SNPs was 6,585,081 and was reduced to 439,775 based on LD pruning with PLINK2. Known clades are shown with colored labels. Although lacking bootstrap support, the North American and Kazakhstan/Georgian sexual populations are two dispersed genetic groups. Clade 1 contains the Ug99 lineage of wheat stem rust and the sister clade to clades 1 and 2 is rye stem rust (*Pgs*; *Puccinia graminis* f. sp. *secalis*), indicating a close relationship between *Pgs* and Clades 1 and 2. A total of 345 isolates of *P. graminis* were included in the analysis. Oat stem rust (*Pga*; *Puccinia graminis* f. sp. *avenae*) is used as an outgroup. Bootstrap support above 80% is shown as red dots at the center point of branches (based on 746 bootstraps with convergence).

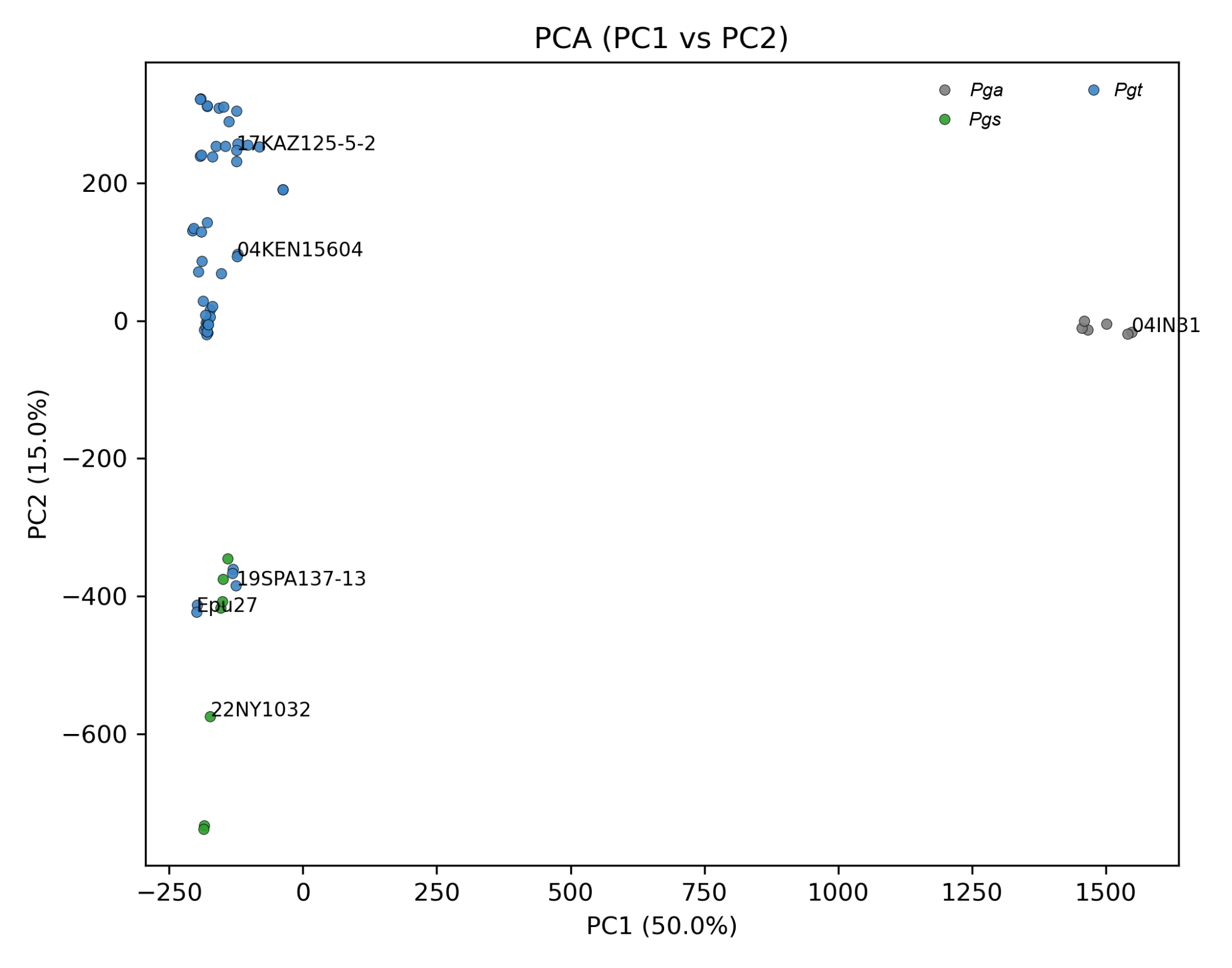

**Figure S3. Principal component analysis of the *Pg* isolates using 9,677,946 SNPs of variants called on the Pgt21-0 genome.**

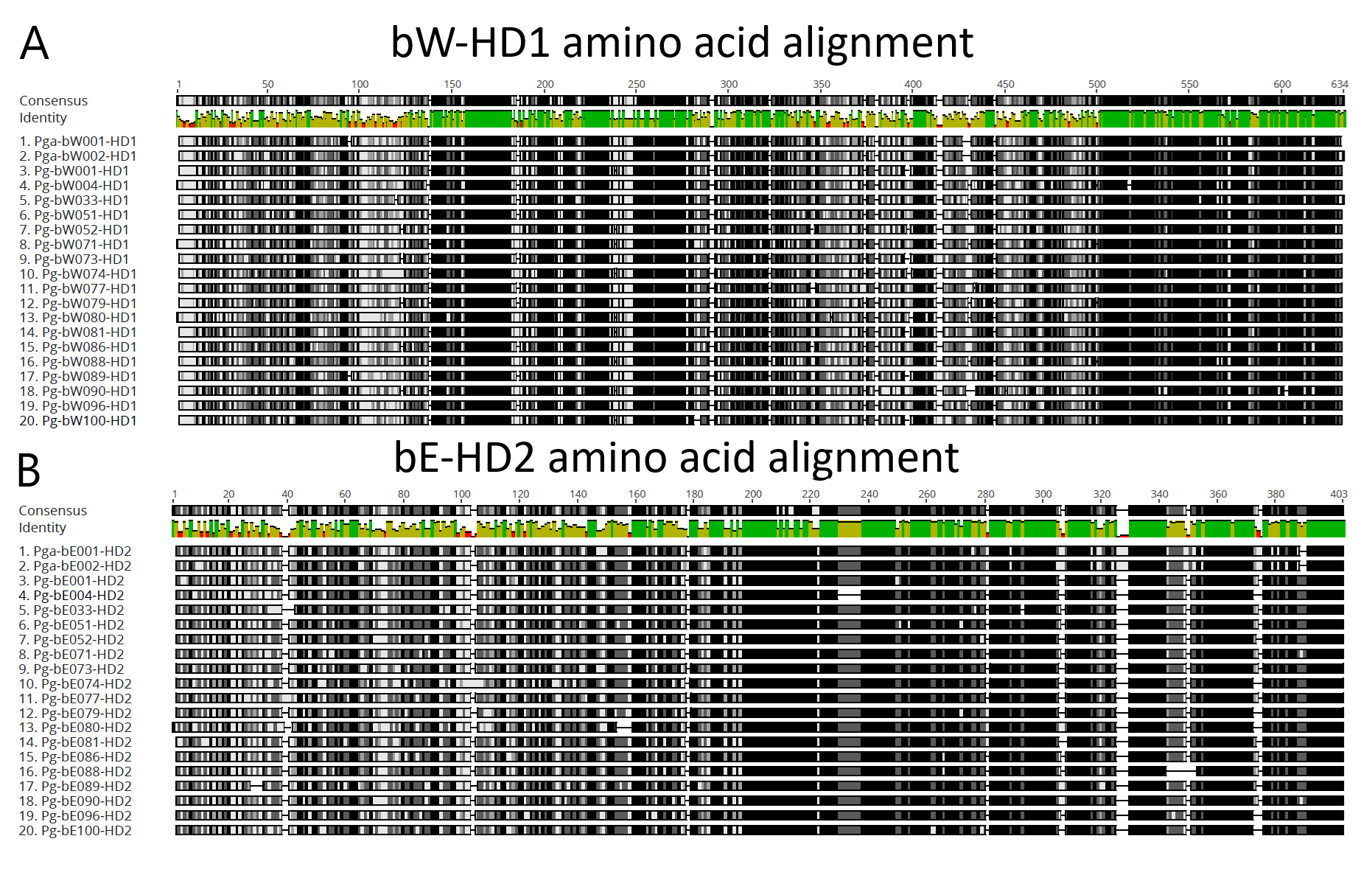

**Figure S4. Amino acid alignments of (A) *Pg* *bW-HD1* (B) and *Pg bE-HD2* genes.** Green regions represent 100% identity, yellow 99-50%, and red 49-0%.

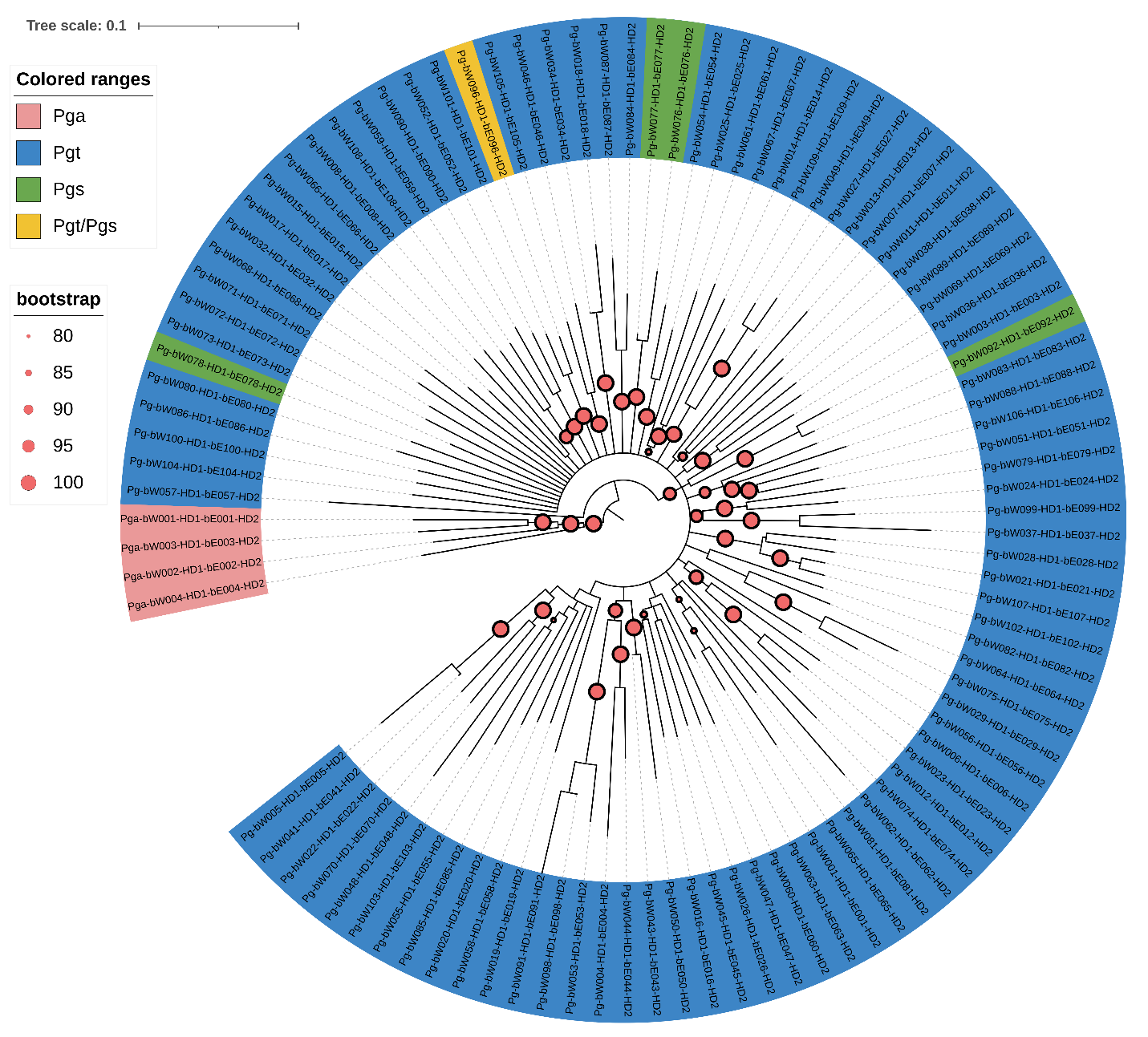

**Figure S5. Neighbor joining tree developed with the Jukes Cantor model based on 1,000 times with bootstrap values >80 reported of 100 *Pg* HD haplotypes.**

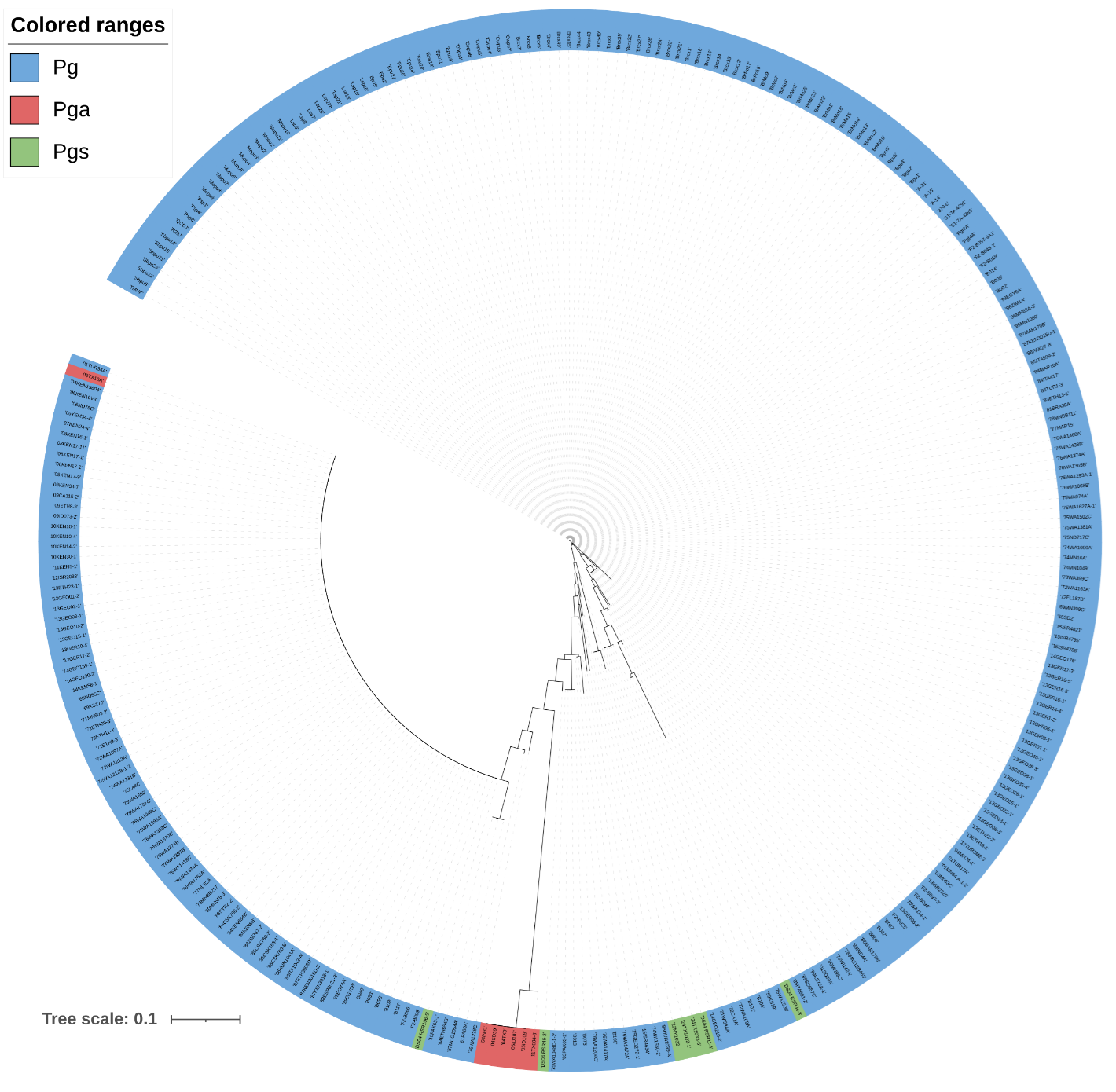

**Figure S6. *Mfa1* neighbor-joining tree developed with the Jukes-Cantor model based on 18 DNA SNPs.**

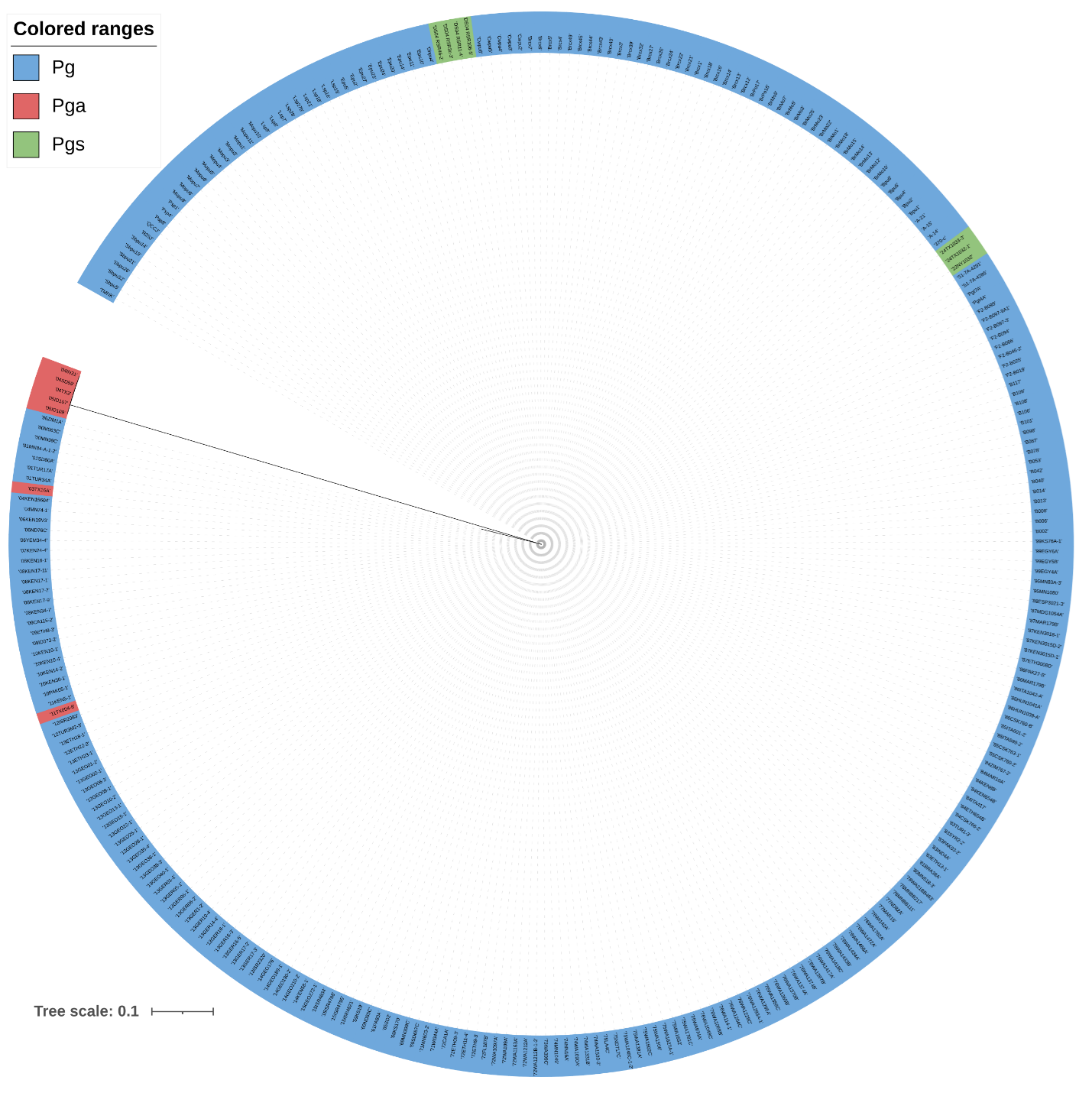

**Figure S7. *Mfa2* neighbor-joining tree developed with the Jukes-Cantor model based on 5 DNA SNPs.**

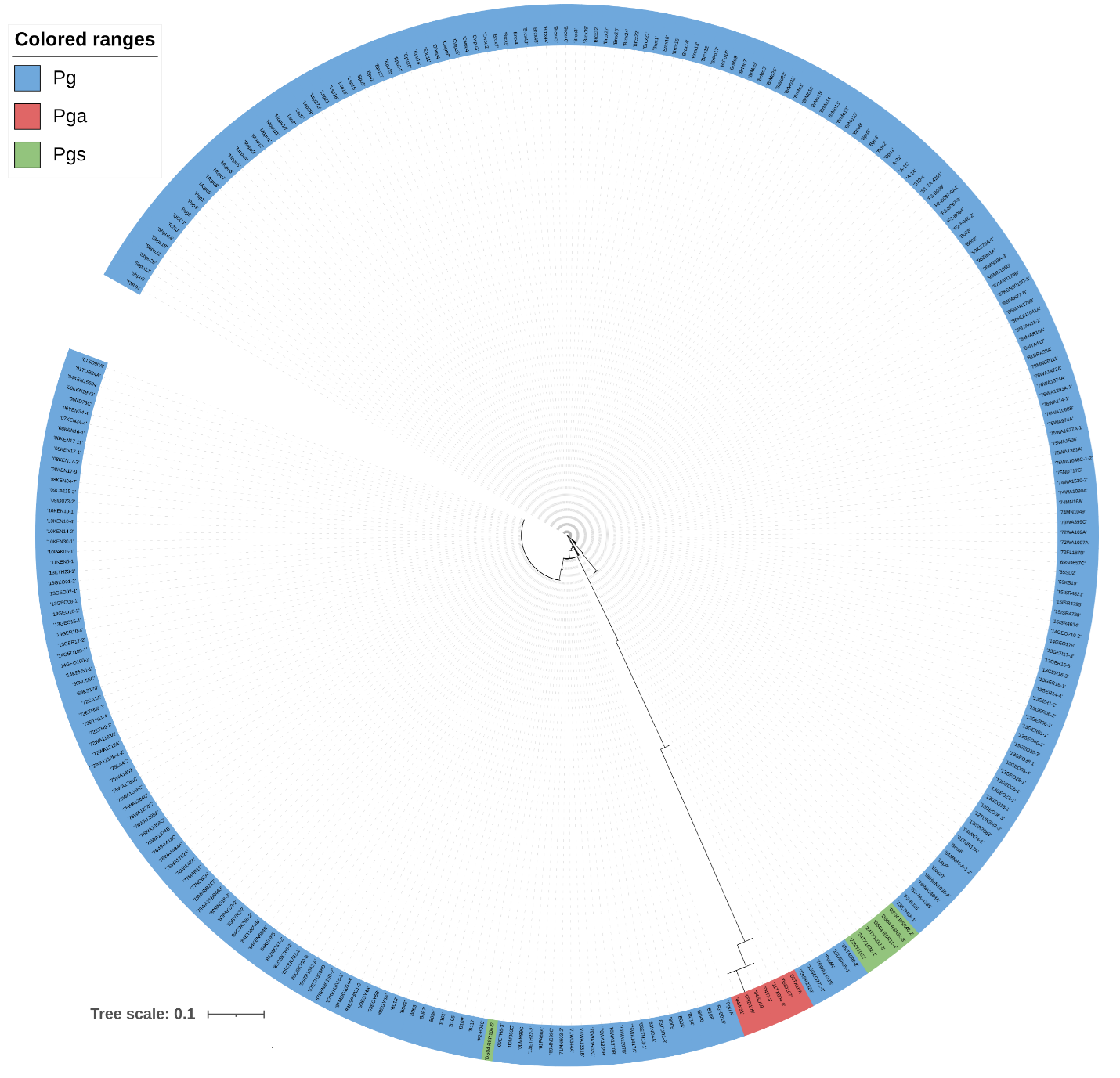

**Figure S8. *Pra* *STE3.2-2* Pgt21-0 neighbor-joining tree developed with the Jukes-Cantor model based on 72 DNA SNPs.**

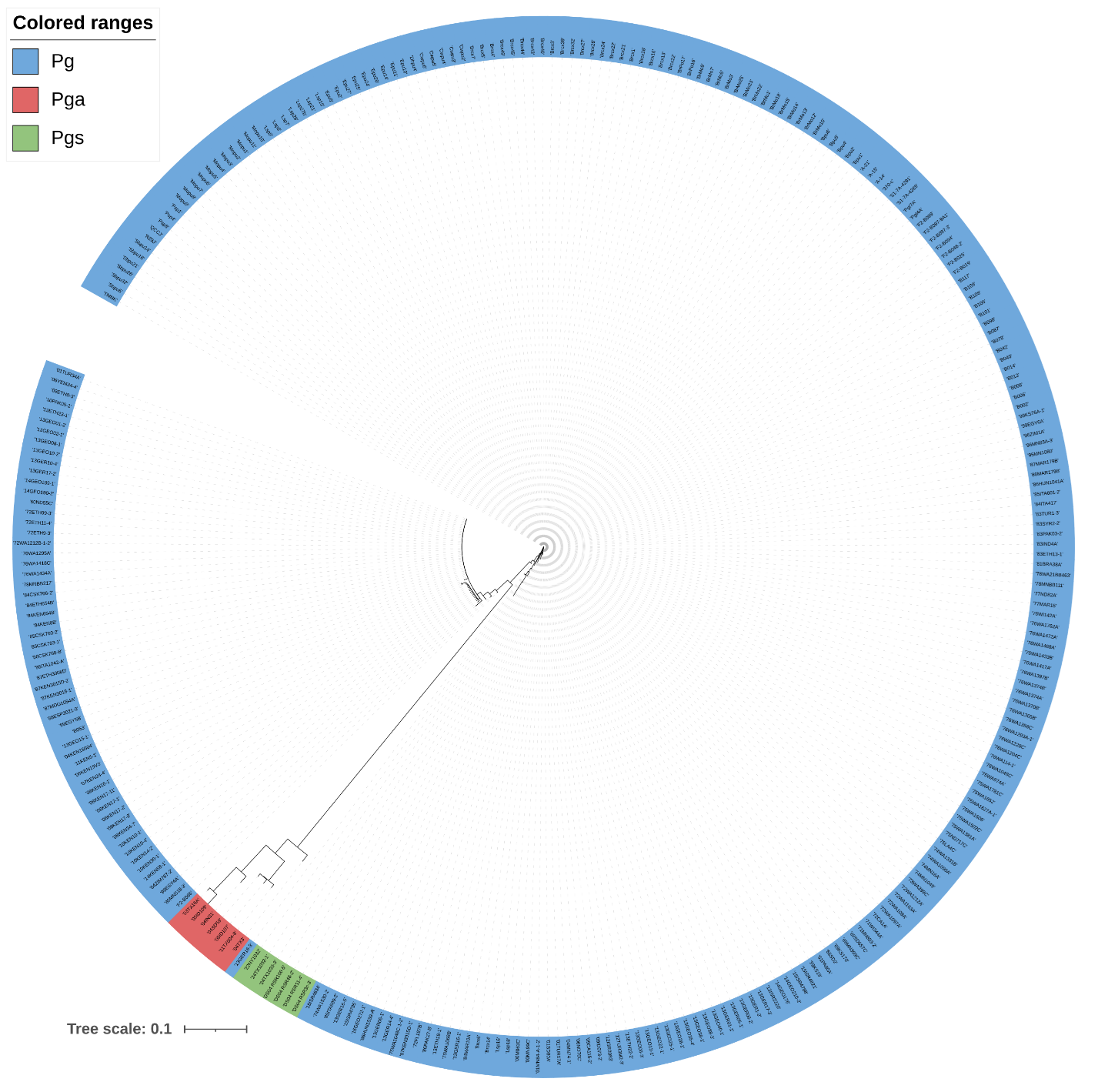

**Figure S9. *Pra* *STE3.2-3* Pgt21-0 neighbor-joining tree developed with the Jukes-Cantor model based on 75 DNA SNPs.**

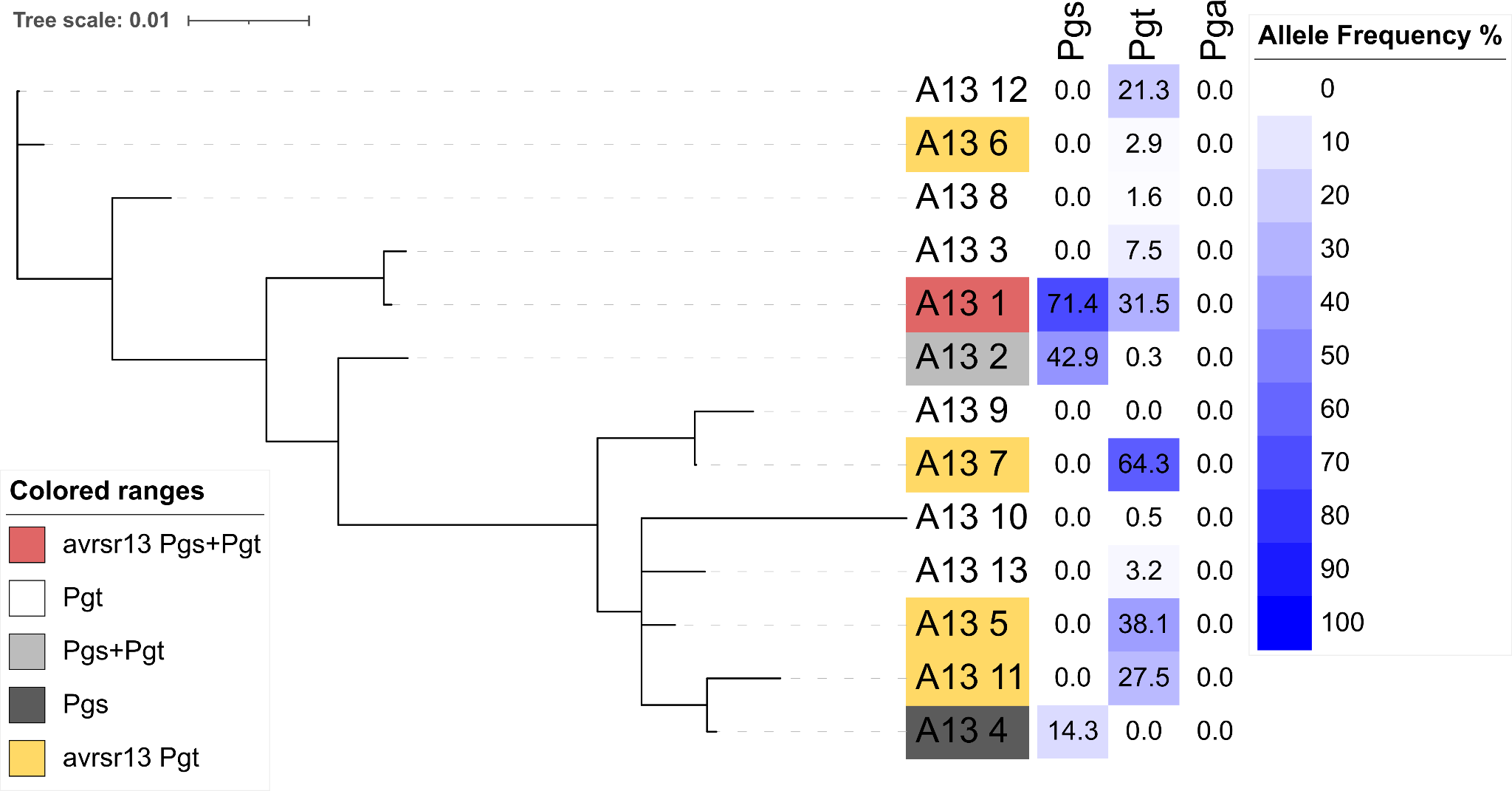

**Figure S10. Phylogenetic tree of *AvrSr13* alleles and postulated compatibility on *Sr13*.** Avirulence and virulence status is based on an evaluation of published literature and allelic variation based on k-mer analysis (Table S5). Host specificity is based on published literature and established differential sets of representative lines for each species (Supplemental Dataset 1). Allele frequency calculated as a percentage of alleles with 100% containment in different *Pg* *formae speciales*.

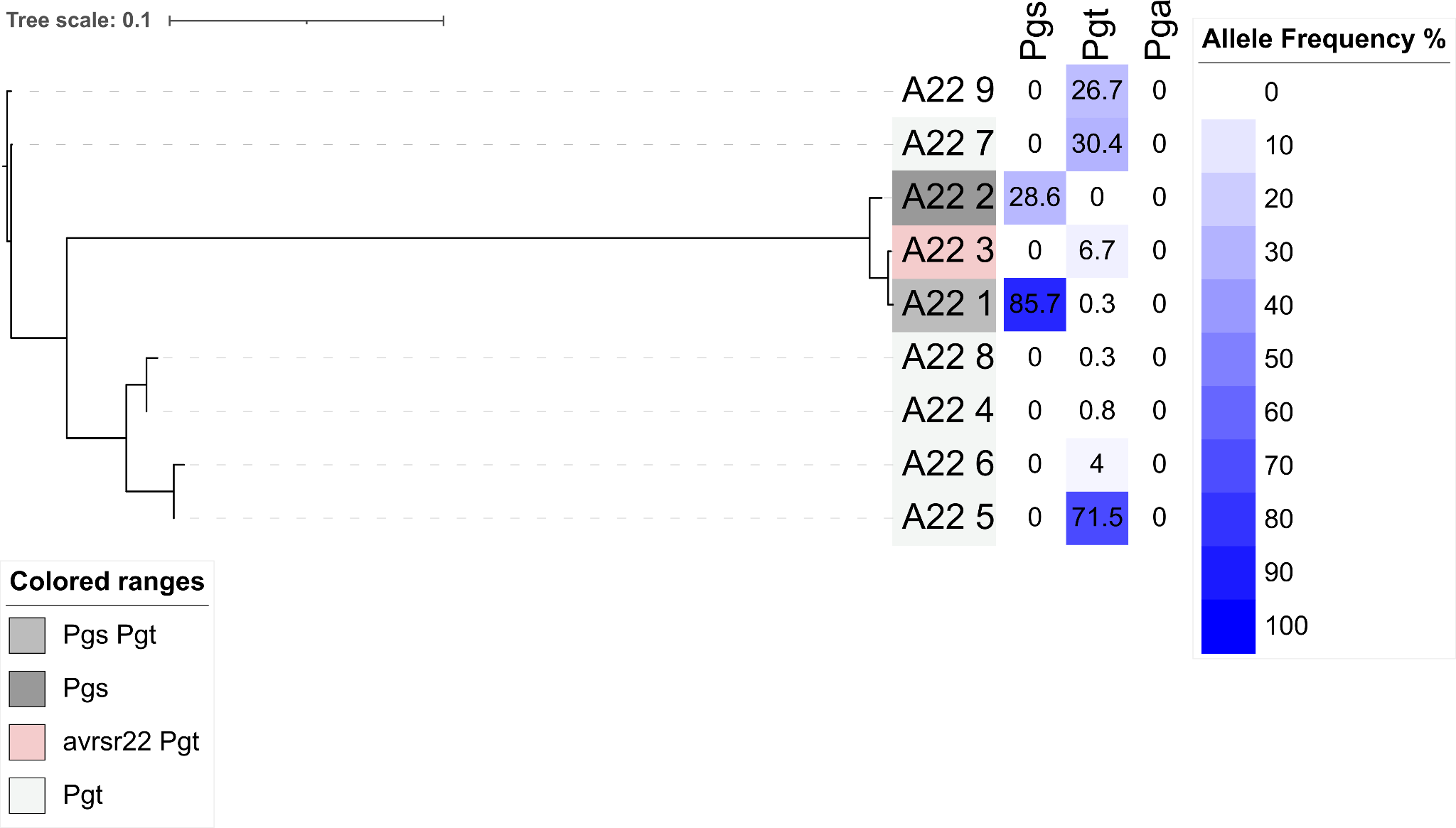

**Figure S11. Phylogenetic tree of *AvrSr22* alleles and postulated compatibility on *Sr22*.** Avirulence and virulence status is based on an evaluation of published literature and allelic variation based on k-mer analysis (Table S5). Host specificity is based on published literature and established differential sets of representative lines for each species (Supplemental Dataset 1). Allele frequency calculated as a percentage of alleles with 100% containment in different *Pg* *formae speciales*.

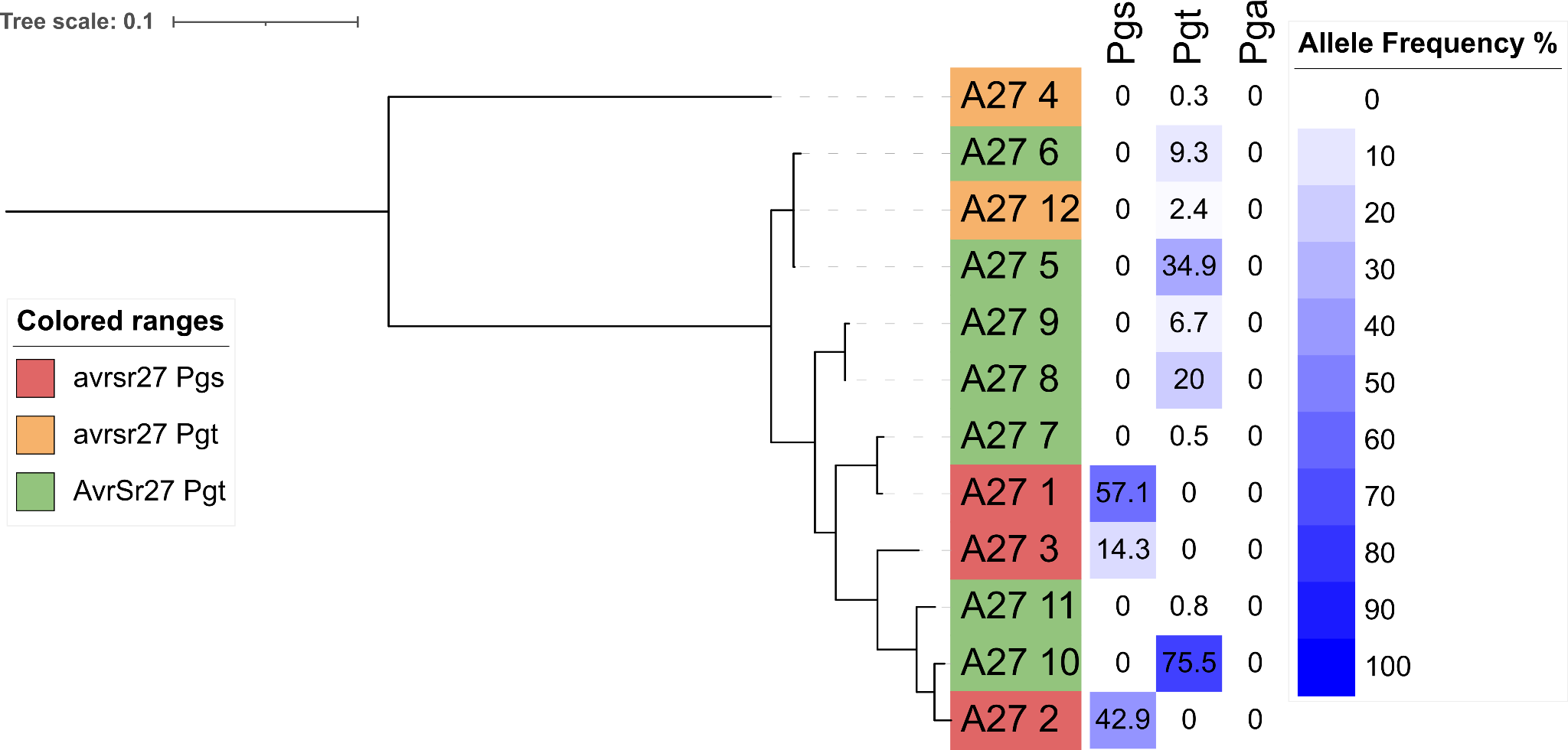

**Figure S12. Phylogenetic tree of *AvrSr27* alleles and postulated compatibility on *Sr27*.** Avirulence and virulence status is based on an evaluation of published literature and allelic variation based on k-mer analysis (Table S5). Host specificity is based on published literature and established differential sets of representative lines for each species (Supplemental Dataset 1). Allele frequency calculated as a percentage of alleles with 100% containment in different *Pg* *formae speciales*.

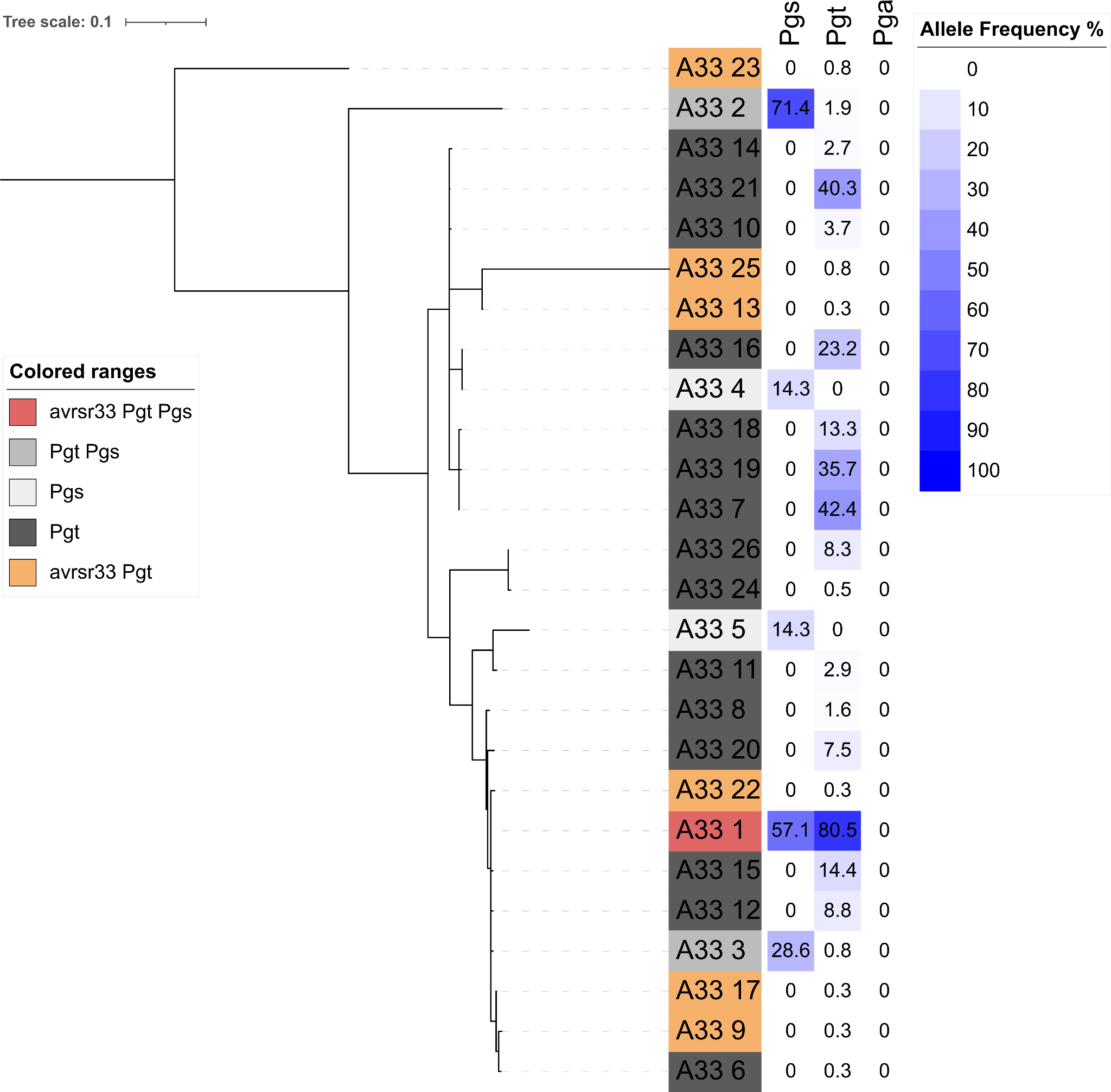

**Figure S13. Phylogenetic tree of *AvrSr33* alleles and postulated compatibility on *Sr33*.** Avirulence and virulence status is based on an evaluation of published literature and allelic variation based on k-mer analysis (Table S5). Host specificity is based on published literature and established differential sets of representative lines for each species (Supplemental Dataset 1). Allele frequency calculated as a percentage of alleles with 100% containment in different *Pg* *formae speciales*.

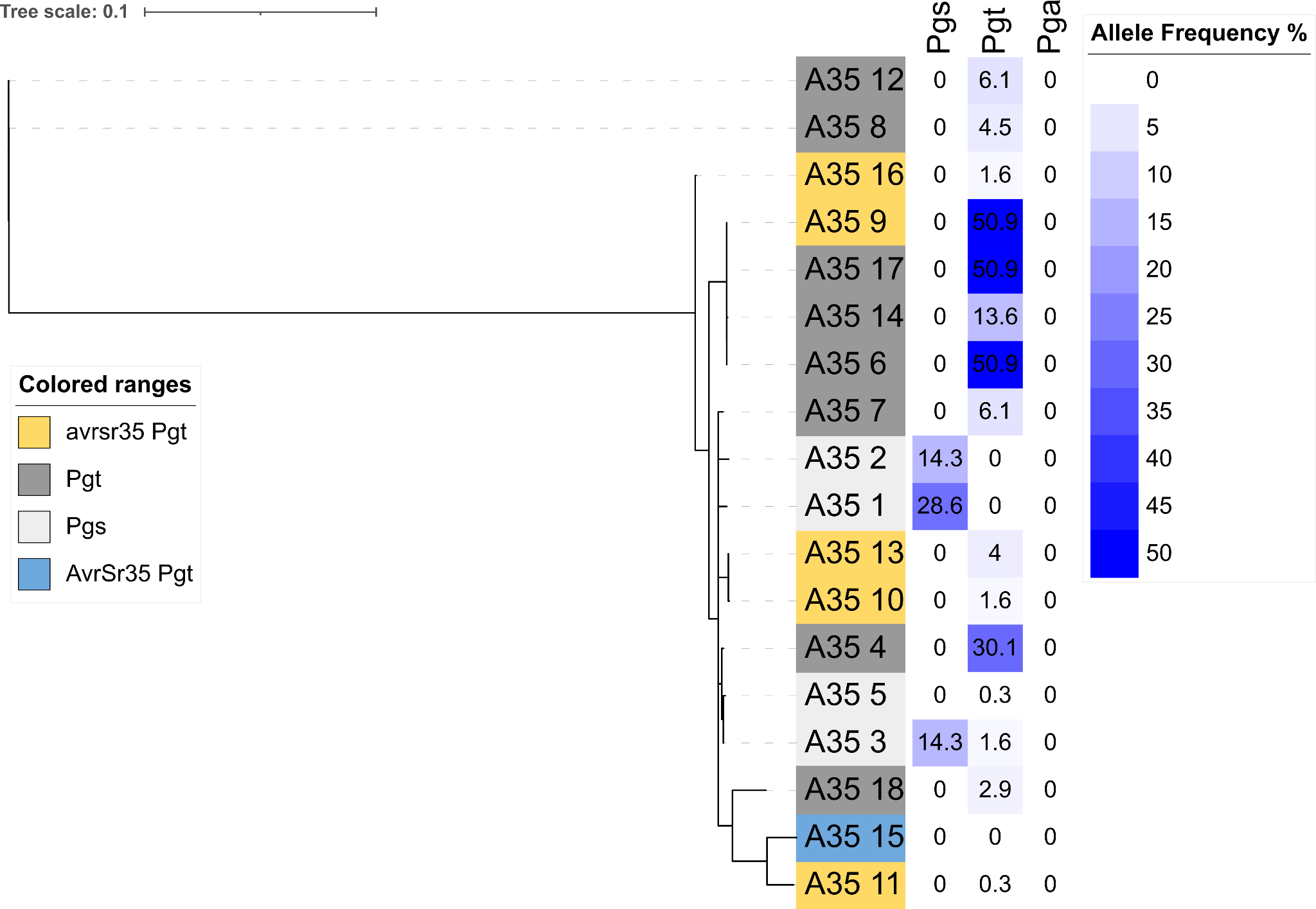

**Figure S14. Phylogenetic tree of *AvrSr35* alleles and postulated compatibility on *Sr35*.** Avirulence and virulence status is based on an evaluation of published literature and allelic variation based on k-mer analysis (Table S5). Host specificity is based on published literature and established differential sets of representative lines for each species (Supplemental Dataset 1). Allele frequency calculated as a percentage of alleles with 100% containment in different *Pg* *formae speciales*.

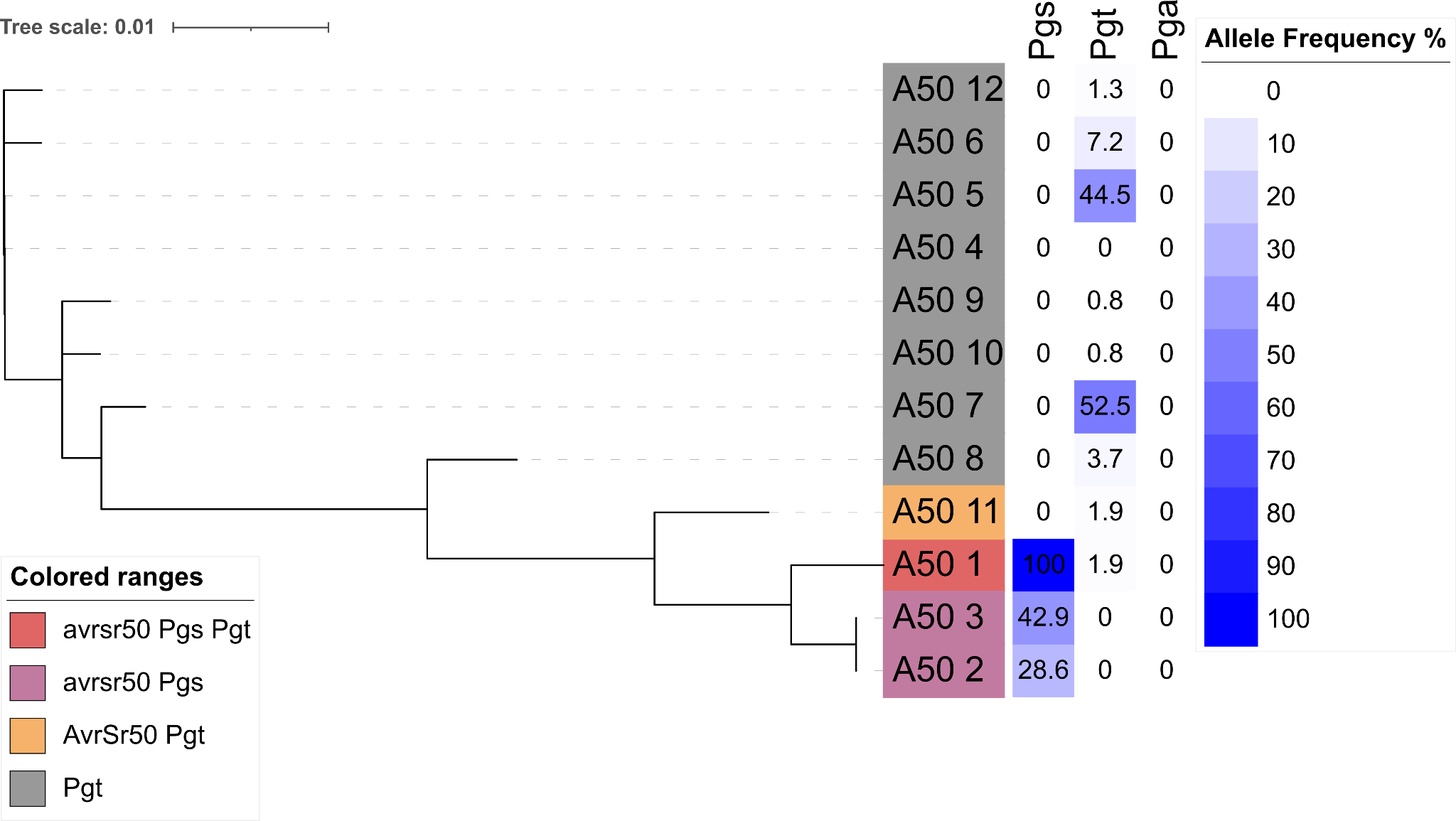

**Figure S15. Phylogenetic tree of *AvrSr50* alleles and postulated compatibility on *Sr50*.** Avirulence and virulence status is based on an evaluation of published literature and allelic variation based on k-mer analysis (Table S5). Host specificity is based on published literature and established differential sets of representative lines for each species (Supplemental Dataset 1). Allele frequency calculated as a percentage of alleles with 100% containment in different *Pg* *formae speciales*.

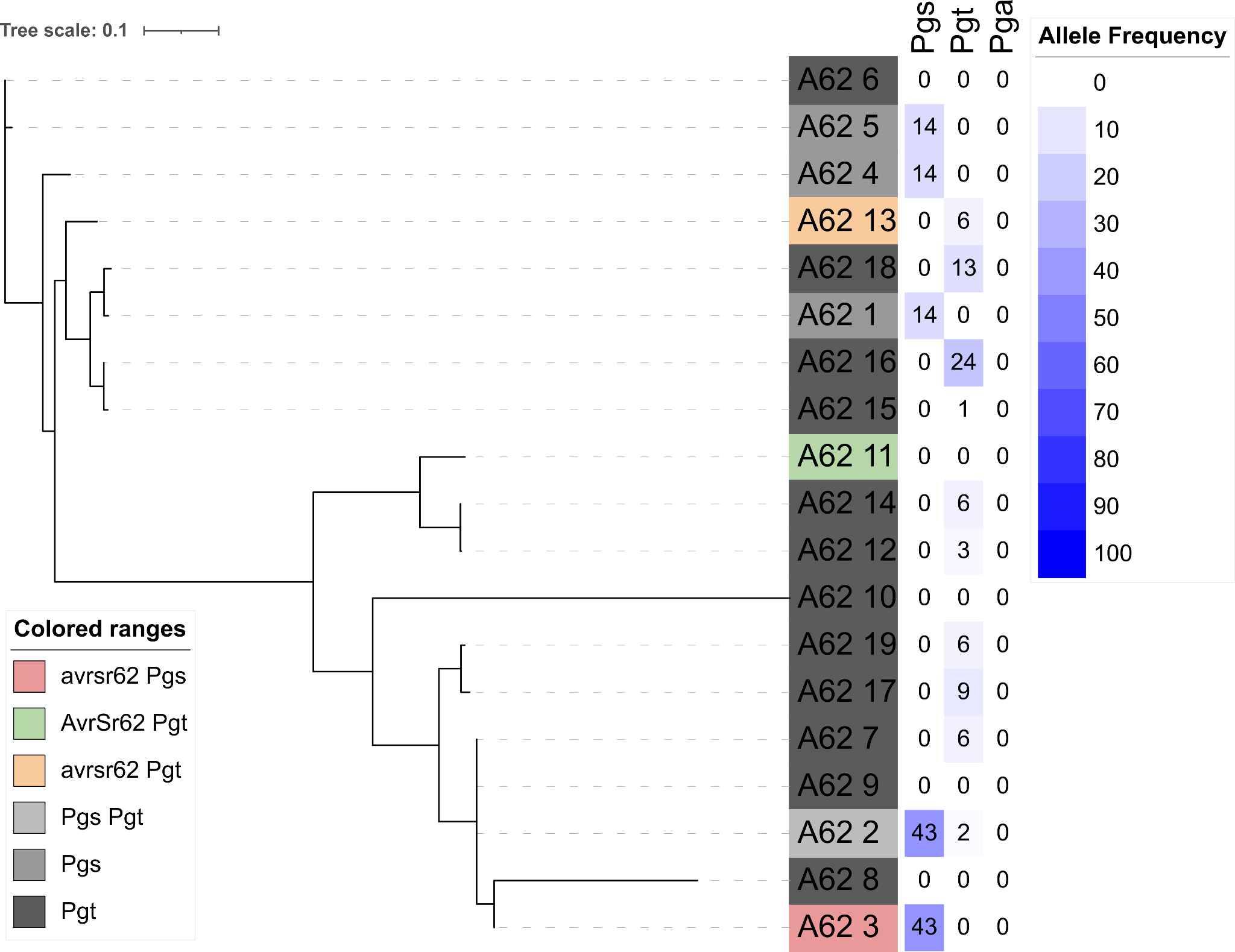

**Figure S16. Phylogenetic tree of *AvrSr62* alleles and postulated compatibility on *Sr62*.** Avirulence and virulence status is based on an evaluation of published literature and allelic variation based on k-mer analysis (Table S5). Host specificity is based on published literature and established differential sets of representative lines for each species (Supplemental Dataset 1). Allele frequency calculated as a percentage of alleles with 100% containment in different *Pg* *formae speciales*.

| **Table S1. HD haplotypes contributing to novel *Pg* races, clades, or epidemics.** | | | |
| --- | --- | --- | --- |
| HD haplotype name | Isolate^a^ | *Pg* f. sp.^b^ | Event |
| Pg-bW008-HD1-bE008-HD2 | 15ISR4788 | *tritici* | TKTTF 'Digalu', TTRTF |
| Pg-bW007-HD1-bE007-HD2 | 14GEO189-1 | *tritici* | TTRTF |
| Pg-bW100-HD1-bE100-HD2 | Epu27 | *tritici* | Novel *Pg* clade |
| Pg-bW074-HD1-bE074-HD2 | 01MN84-A-1-2 | *tritici* | TTTTF, RCRSC |
| Pg-bW051-HD1-bE051-HD2 | 17KAZ108-3 | *tritici* | *Sr32*, *Sr40* novel virulence |
| Pg-bW052-HD1-bE052-HD2 | 17KAZ108-3 | *tritici* | *Sr32*, *Sr40* novel virulence |
| Pg-bW004-HD1-bE004-HD2 | Ug99 | *tritici* | Ug99 epidemic |
| Pg-bW003-HD1-bE097-HD2 | Ug99 | *tritici* | Ug99 epidemic |
| Pg-bW104-HD1-bE104-HD2 | 19SPA124-9-1 | *tritici* | *Sr31* virulence |
| Pg-bW105-HD1-bE105-HD2 | 19SPA124-9-1 | *tritici* | *Sr31* virulence |
| Pg-bW106-HD1-bE106-HD2 | 19SPA127-10-1 | *tritici* | *Sr31* virulence |
| Pg-bW108-HD1-bE108-HD2 | 19SPA127-10-1 | *tritici* | *Sr31* virulence |
| Pg-bW107-HD1-bE107-HD2 | 19SPA137-13 | *tritici* | *Sr31* virulence |
| Pg-bW109-HD1-bE109-HD2 | 19SPA137-13 | *tritici* | *Sr31* virulence |
| Pg-bW096-HD1-bE096-HD2 | Mopu1 | *tritici, secalis* | 1970's Washington State |
| ^a^Isolate assembly from which the HD haplotype was discovered. | | | |
| ^b^Classification of haplotype of *P. graminis forma specialis* based on virulence on rye or wheat. | | | |
